# How a Hemicarcerand Incarcerates Guests at Room Temperature Decoded with Modular Simulations

**DOI:** 10.1101/2020.11.20.390328

**Authors:** Katherine G. McFerrin, Yuan-Ping Pang

## Abstract

Hemicarcerands are host molecules created to study constrictive binding with guest molecules for insights into the rules of molecular complexation. However, the molecular dynamics simulations that facilitate such studies have been limited because three-dimensional models of hemicarcerands are tedious to build and their atomic charges are complicated to derive. There have been no molecular dynamics simulations of the reported water-soluble hemicarcerand (Octacid4) that explain how it uniquely encapsulates its guests at 298 K and keeps them encapsulated at 298 K in NMR experiments. Herein we report a modular approach to hemicarcerand simulations that simplifies the model building and charge derivation in a manner reminiscent of the approach to protein simulations with truncated amino acids as building blocks. We also report that *apo* Octacid4 in water adopts two clusters of conformations, one of which has an equatorial portal open thus allowing guests to enter the cavity of Octacid4, in microsecond molecular dynamics simulations performed using the modular approach at 298 K. Under the same simulation conditions, the guest-bound Octacid4 adopts one cluster of conformations with all equatorial portals closed thus keeping the guests incarcerated. These results explain the unique constrictive binding of Octacid4 and suggest that the guest-induced host conformational change that impedes decomplexation is a previously unrecognized conformational characteristic that promotes strong molecular complexation.

## Introduction

Hemicarcerands comprise two identical bowl-shaped resorcinarene fragments tethered with four linkers (Fig. 1). These molecules, containing four equatorial portals in the linker region and two axial portals in the resorcinarene region, are developed as hosts to encapsulate small-molecule guests in a typical way that the guests enter and exit the host cavity at high temperatures and remain in the cavity at low temperatures^1,2^. The binding characteristics and solubility of hemicarcerands are governed mostly by the linker structures. Octaacid 4 (Octacid4; tethered with a 4,6-dimethylisophthalic acid fragment as shown in Fig. 1) is reportedly a unique hemicarcerand that is water soluble and can form host•guest complexes (hemicarceplexes) with small molecules in water without the need to raise temperature^3^. Therefore, the Octacid4-like hemicarceplexes are useful model systems for the study of constrictive binding, which is a type of molecular complexation that is affected by the activation energy required for a guest to enter the host cavity through a size-restricting portal^4–6^, to obtain insights into the rules of molecular complexation.

**Fig. 1.**
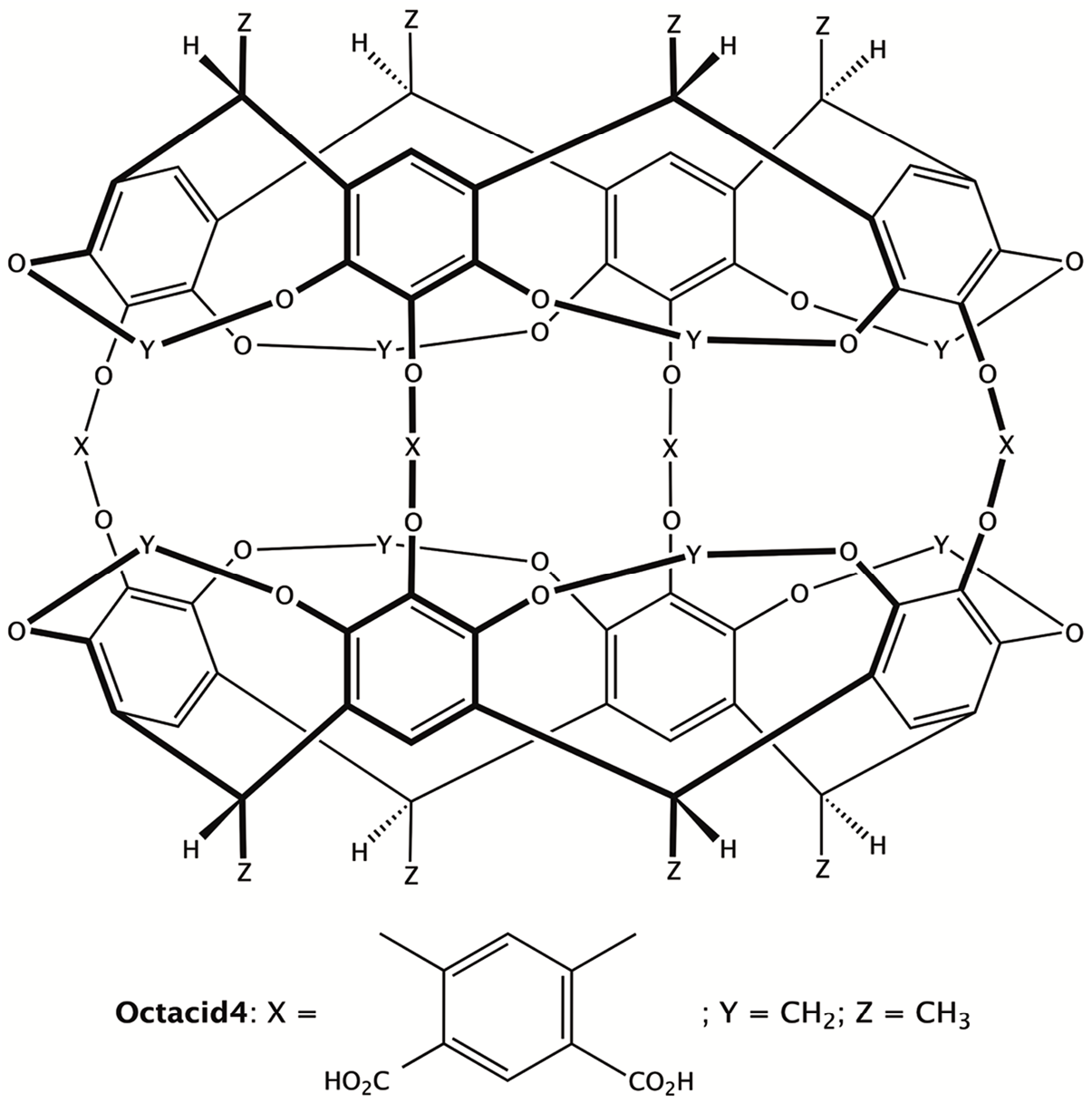
Chemical structures of a generic hemicarcerand and Octacid4.

To date, only a few molecular dynamics (MD) simulations of hemicarcerands or hemicarceplexes have been reported for the facilitation of constructive bindings studies^7–10^. This scarcity is partly because hemicarcerands have more than 200 atoms, making their three-dimensional models tedious to build. The scarcity is also due to the complexity of hemicarcerands, all of which have multiple conformations^8,11^. As shown in Fig. 2, variations of the torsions along the four linkers of Octacid4 can result in many distinct conformations, which complicates the derivation of the conformation-dependent atomic charges of the hemicarcerand. These technical complexities, as detailed below, may explain why there have hitherto been no aqueous MD simulations of Octacid4 to understand how it uniquely encapsulates various small-molecule guests at 298 K and keeps them encapsulated at the same temperature such that the bound guests can be differentiated in their NMR spectra from those in the bulk phase^3^.

**Fig. 2.**
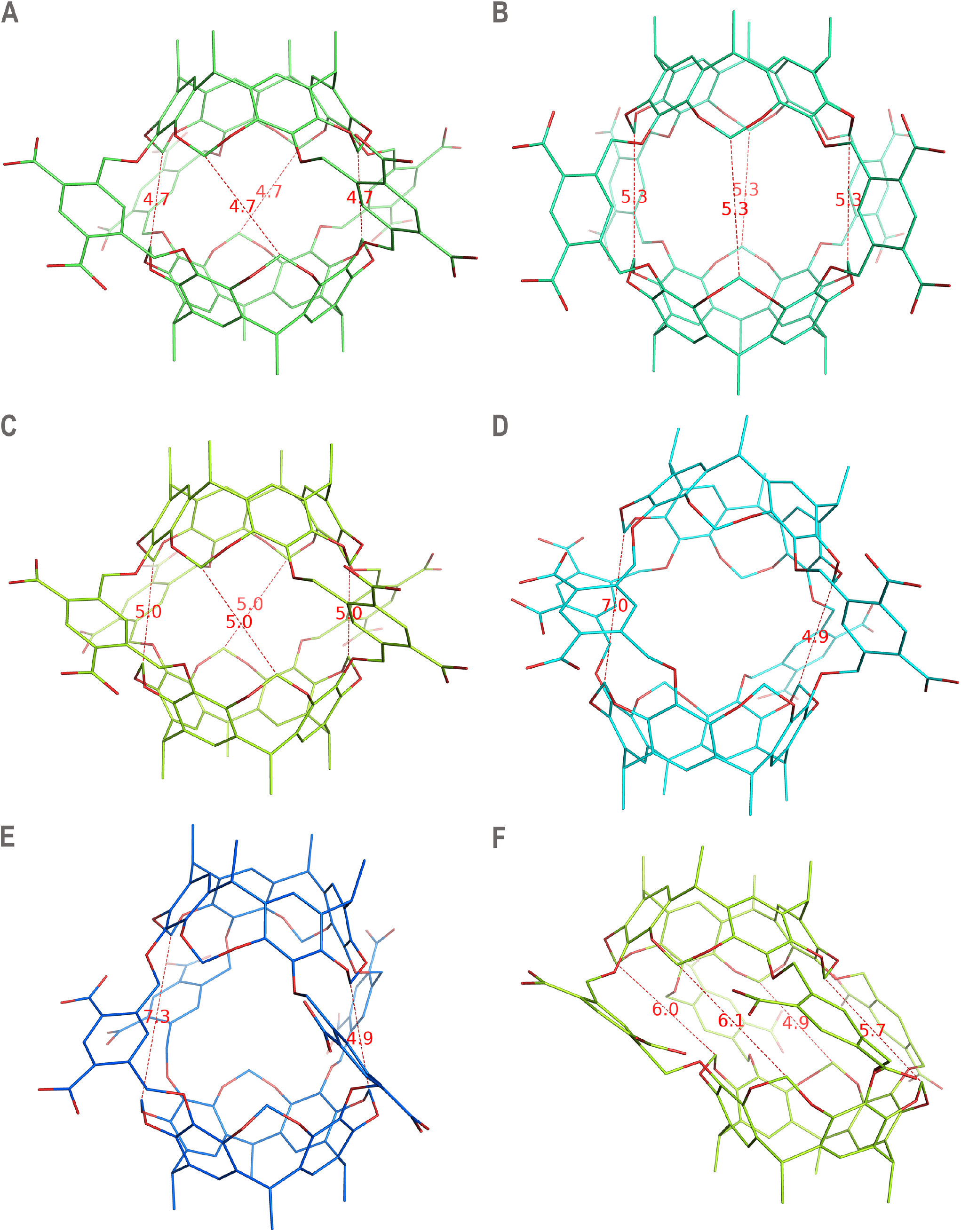
Distinct conformations of Octacid4. Distances shown by thin dashed lines to indicate the open or closed equatorial portal are in Angstroms. Oxygen atoms are in red. Carbon atoms are in lime, green cyan, limon, cyan, marine, or yellow orange. Counter ions and water molecules of aqueous Octacid4 are not displayed for clarity. **(A)** Energy-minimization– derived Octacid4 in vacuo with all equatorial portals closed. **(B)** Energy-minimization– derived Octacid4 in vacuo with all equatorial portals open. **(C)** The most populated Octacid4 conformation with all equatorial portals closed in the aqueous MD simulations. **(D)** The second most populated Octacid4 conformation with one equatorial portal open in the aqueous MD simulations. **(E)** An Octacid4 conformation with one equatorial portal open in the aqueous MD simulations**. (F)** Another Octacid4 conformation with equatorial portals open in the aqueous MD simulations.

Herein we report a modular method that simplifies the model building and charge derivation for MD simulations of Octacid4 and its hemicarceplexes. We also report the characterization of the host cavity and motions of the encapsulated guests that explains the unique constrictive binding of Octacid4 and offer new mechanistic insight into molecular complexation.

## Results

### The modular approach

In addition to the general technical complexities in hemicarcerand simulations noted above, the charge derivation for water-soluble hemicarcerands such as Octacid4 is particularly complicated. Because of the need to balance the atomic charges of the water-soluble hemicarcerand with the charges of the aqueous solvent and the charges of the small-molecule guest when using an AMBER forcefield such as FF12MClm^12^, the hemicarcerand charges need to be derived from ab initio calculations using the HF/6-31G* basis set that uniformly overestimates the polarity of the molecule. This is because (1) aqueous solvent models (such as the widely used TIP3P empirical water model) include polarization effects due to the empirical calibration to reproduce the density and enthalpy of liquid vaporization^13^, and (2) the small-molecule guest bears the restrained electrostatic potential (RESP) charges that are derived from ab initio calculations using the HF/6-31G* basis set^14–16^. Further, to obtain the atomic charges of a water-soluble hemicarcerand without any bias toward one particular conformation of the molecule, the hemicarcerand charges need to be obtained from ab initio calculations at the HF/6-31G*//HF/6-31G* level with multiple conformations of the molecule followed by using the Lagrange multiplier to force identical charges on equivalent atoms in these conformations^16^. These ab initio calculations and the Lagrange multiplier are computation-demanding and labor-intensive, respectively. For example, the ab initio calculation of one Octacid4 conformation at the HF/6-31G*//HF/6-31G* level took ~168 CPU hours using the computers at the University of Illinois Urbana-Champaign National Center for Supercomputing Applications.

During the course of our Octacid4 simulations, we recognized that the technical complexities described above differ little from those of MD simulations of proteins with large numbers of atoms and conformations. One known approach to circumventing the complexities of protein MD simulations is to perform the simulations using the modular concept underlying the second-generation AMBER forcefield^17^. This forcefield builds a protein model with truncated amino acids as building blocks and derives the atomic charges of each building block from its multiple conformations using the Lagrange multiplier to force identical charges on equivalent atoms in these conformations^16^. Given the effectiveness of the modular concept as demonstrated by the reported autonomous miniprotein folding MD simulations that achieved agreements between simulated and experimental folding times within factors of 0.6–1.4^18^, we used to the modular concept to simplify the model building and charge derivation for hemicarcerands and devised a modular method for hemicarcerand simulations. We describe below two key attributes of our method for performing the MD simulations of Octacid4.

### The building block of Octacid4

Because Octacid4, similar to most hemicarcerands, has C4 symmetry, we divided it into four identical building blocks (termed HC1; Fig. 3). In this building block, C1 and C20 are designated as the head and tail atoms of the residue, respectively, similar to the designation of head and tail atoms in a truncated amino acid residue for protein simulations. Topology-wise, we built Octacid4 as shown in Fig. 3 with a sequence of HC1-HC1-HC1-HC1.

**Fig. 3.**
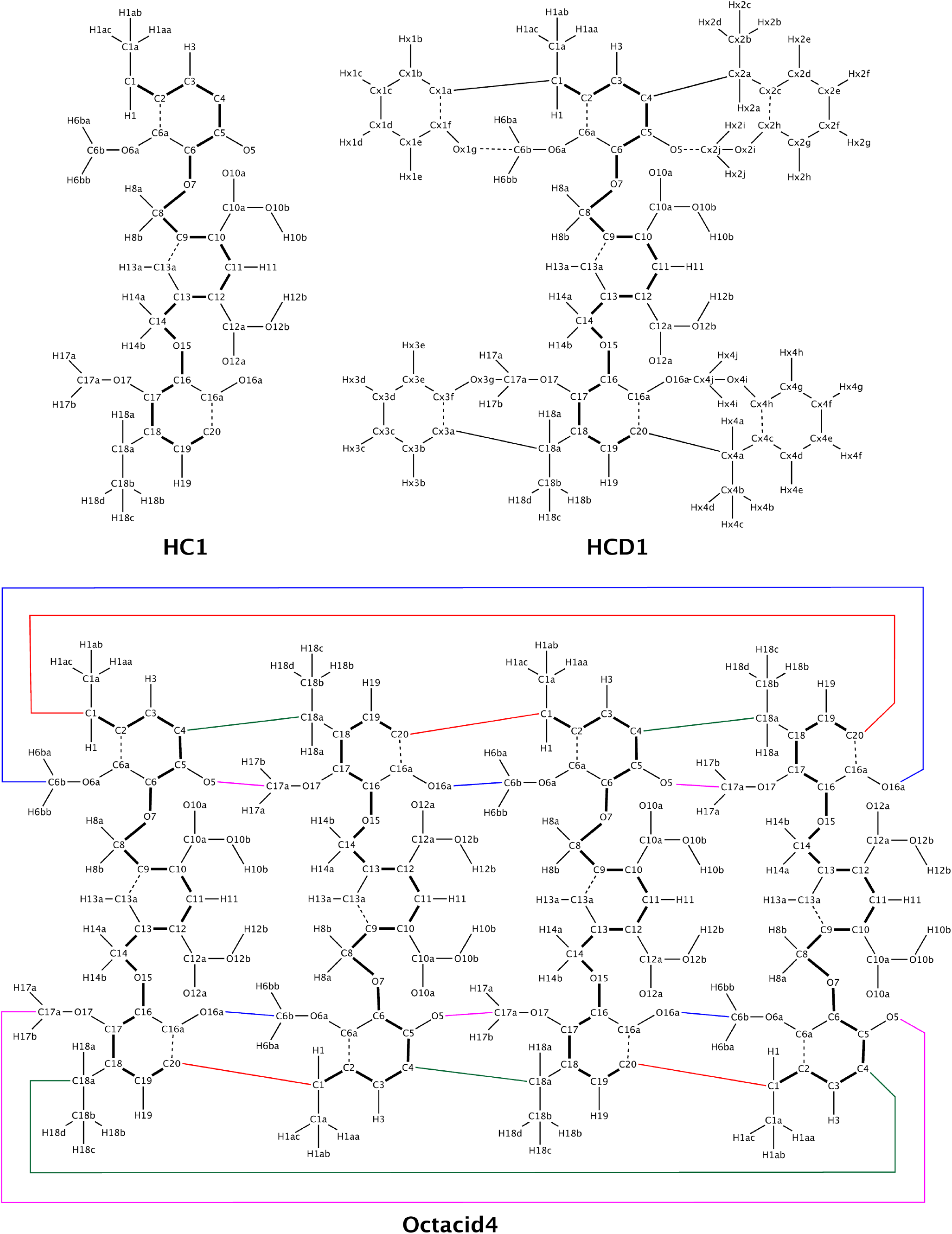
Chemical structures of HC1 and HCD1 and the assembly of HC1 into Octacid4. Thick lines indicate the bonds between main-chain atoms. Thin lines indicate the bonds between side-chain atoms or between a main-chain atom and a side-chain atom. Thin dashed lines and colored lines indicate the intra- and inter-residue bonds, respectively.

First, we assembled the four residues into a linear molecule. This assembly followed the same scheme as that for assembling protein residues—namely, constructing a covalent bond between the tail atom of the preceding residue and the head atom of the following residue. Second, we converted the linear molecule to a cyclic molecule by specifying a cross-link (*viz*., constructing a covalent bond) between the head atom of the first HC1 and the tail atom of the last HC1. Third, we converted the cyclic molecule to Octacid4 by specifying (*i*) four cross-links between C4 of the preceding HC1 and C18a of the following HC1, (*ii*) four cross-links between O5 of the preceding HC1 and C17a of the following HC1, and (*iii*) four cross-links between O16a of the preceding HC1 and C6b of the following HC1.

The Octacid4 sequence and all cross-links can be specified all together, thus making the hemicarcerand model building conceptually simple, algorithmic, and generalizable. Notably the HC1 building block can, in the same manner, be further divided into three sub-building blocks for hemicarcerands without C4 symmetry.

### Atomic charges of HC1

Given the HC1 residue defined above, we obtained the atomic charges of Octacid4 via the RESP charge derivation for HC1 with the following specific conditions, in addition to the general conditions that are the same as those for protein RESP charges (such as the two-stage fitting for the methyl and methylene groups and forcing identical charges on equivalent intramolecular atoms)^16,17^.

First, we converted HC1 to HCD1 by attaching blocking groups to the junction atoms of HC1 (Fig. 3) as these groups are needed to mimic the polar and aromatic groups abutting the junction atoms in Octacid4. We then performed the ab initio calculation of HCD1 to derive the HCD1 charges by using the Lagrange multiplier to force (1) HCD1 to have a net charge of –2 and (2) the total charge of all blocking groups to be zero, similar to the protein charge derivation from ab initio calculations of the acetyl- and *N*-methyl-blocked amino acids whose blocking group are used to mimic adjacent residues of the central amino acid^16^. Second, to balance the hemicarcerand charges with those of the solvent and the guest, we obtained the HCD1 RESP charges from ab initio calculations of HCD1 at the HF/6-31G*//HF/6-31G* level. Third, to avoid any bias toward one particular conformation of Octacid4, we obtained the HCD1 RESP charges from ab initio calculations using (1) two HCD1 conformations taken from the representative conformations of Octacid4 in vacuo with all equatorial portals open (abbreviated as the open conformation; Fig. 2A) and all equatorial portals closed (abbreviated as the closed conformation; Fig. 2B) and (2) the Lagrange multiplier to force identical charges on equivalent atoms in the two conformations akin to the charge derivation method for proteins that uses the alpha helical and beta strand side-chain conformations^16^. Last, we extracted the HC1 charges from the HCD1 charges.

In contrast to the computation-demanding and labor-intensive charge derivation for the intact Octacid4 molecule, the HC1 charge derivation can be readily performed with HCD1. For example, the ab initio calculation of each of the two HCD1 conformations at the HF/6-31G*//HF/6-31G* level took 31–40 CPU hours, considerably less than the ~168 CPU hours for Octacid4 noted above.

### Characterization of the Octacid4 cavity

Using the modular method described above, we performed 16 sets of 20 316-ns distinct, independent, unrestricted, unbiased, and classical isobaric–isothermal MD simulations of Octacid4 and its hemicarceplexes with seven small-molecule guests at 298 K and 340 K. These simulations were performed with statistical relevance to investigate the change in the cavity volume of *apo* and guest-bound Octacid4 in water. A set of 20 simulations for each guest had an aggregated simulation time of 6.32 μs. The seven guests were dimethyl sulfoxide (DMSO), ethyl acetate (EtOAc), dimethyl acetamide (DMA), 1,4-dioxane, diethylammonium (DEA), *p*-xylene, and naphthalene. These molecules were selected according to their calculated maximal dimensions and molar volumes (Table 1) from the 14 reported guests that formed hemicarceplexes with Octacid4^3^.

**TABLE 1.** Static and dynamic properties of Octacid4 and its complexes.

| Structure | MDG<br>(Å) | GMV<br>(cm<sup>3</sup>/mol) | Temp<br>(K) | Avg Rg (Å)<br>Mean ± SE | SAGS | MoGIO |
| --- | --- | --- | --- | --- | --- | --- |
| Open Apo Octacid4 (vacuo) | — | — | — | 5.86 | — | — |
| Closed Apo Octacid4 (vacuo) | — | — | — | 5.37 | — | — |
| Apo Octacid4 (water) | — | — | 298 | 5.44 ± 0.02 | — | — |
| | | | 340 | 5.45 ± 0.02 | — | — |
| DMSO-Octacid4 (water) | 4.5 | 49 | 298 | 5.41 ± 0.02 | Ball | Free spin |
| | | | 340 | 5.41 ± 0.02 | | |
| 1,4-Dioxane-Octacid4 (water) | 4.8 | 74 | 298 | 5.37 ± 0.01 | Ball | Free spin |
| | | | 340 | 5.38 ± 0.01 | | |
| DMA-Octacid4 (water) | 5.3 | 73 | 298 | 5.40 ± 0.01 | Short rod | Partially free spin |
| | | | 340 | 5.41 ± 0.01 | | |
| EtOAc-Octacid4 (water) | 6.5 | 71 | 298 | 5.40 ± 0.01 | Short rod | Partially free spin |
| | | | 340 | 5.41 ± 0.01 | | |
| DEA-Octacid4 (water) | 6.8 | 72 | 298 | 5.64 ± 0.02 | Long rod | Axial spin |
| | | | 340 | 5.65 ± 0.02 | | |
| *p*-Xylene-Octacid4 (water) | 6.9 | 82 | 298 | 5.56 ± 0.02 | Long rod | Axial spin |
| | | | 340 | 5.59 ± 0.02 | | |
| Naphthalene-Octacid4 (water) | 7.1 | 115 | 298 | 5.65 ± 0.02 | Long rod | Axial spin |
| | | | 340 | 5.66 ± 0.02 | | |

MDG: The maximal dimension of the guest calculated using Gaussian 98. GMV: Guest molar volume calculated using Gaussian 98. Temp: Temperature. Avg Rg: Average radius of gyration from 20 independent and distinct simulations. SAGS: The shape of the average guest structure. MoGIO: The motion of the guest incarcerated in Octacid4. Axial spin: rotation around the axial axis. Free spin: rotations evenly around multiple axes. Partially free spin: rotations around one axis more frequently than around the orthogonal axis.

As apparent in Fig. 4, the Octacid4 cavity is confined by a set of atoms shown with the sphere model. Therefore, we used the variation in the radius of gyration (Rg) of these atoms to estimate the change in cavity volume. Overall, the standard errors for the average Rgs of the cavity for the 16 sets of MD simulations were either 0.01 Å or 0.02 Å (Table 1), demonstrating the convergence for each simulation of the 16 sets and the statistical rigor of these simulations.

**Fig. 4.**
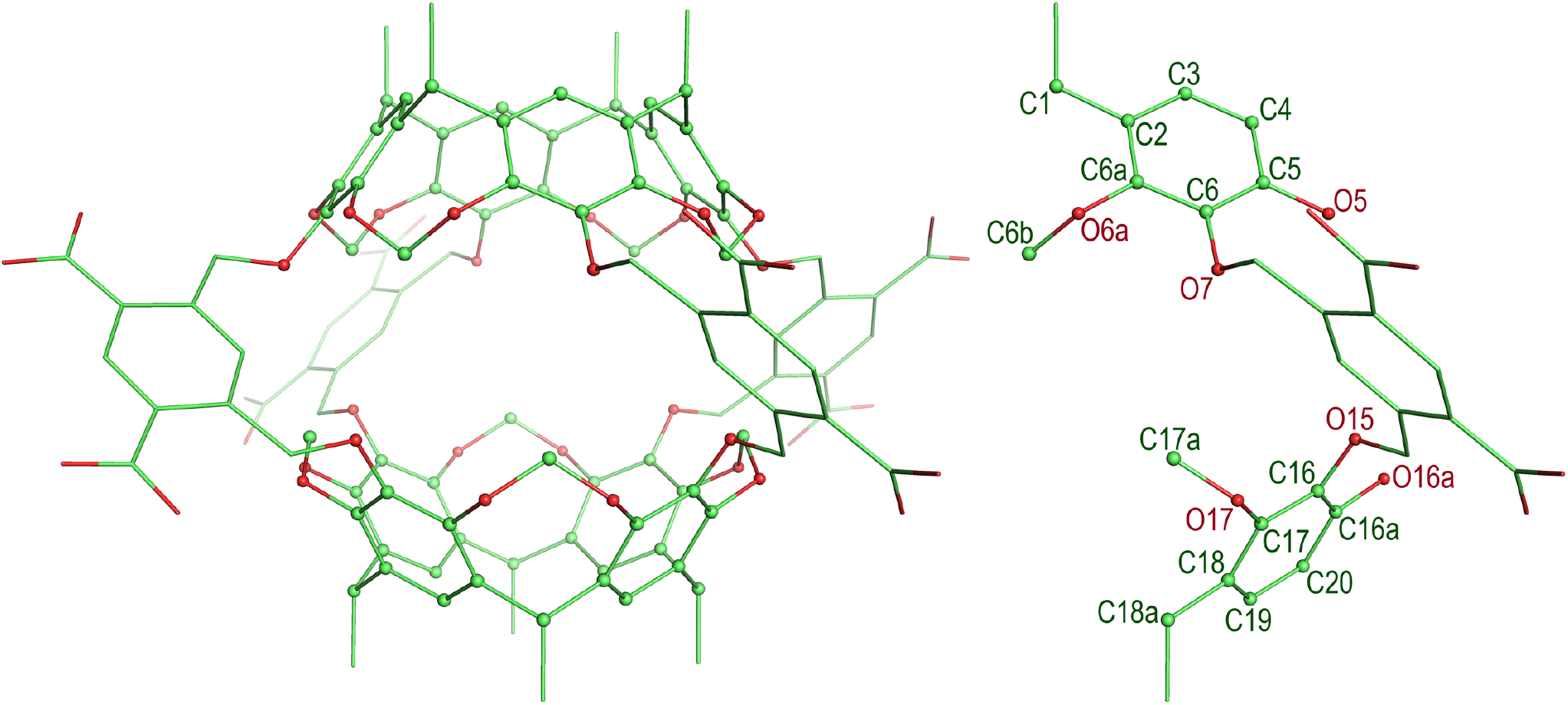
Labels of the atoms in the HC1 residue that confine the Octacid4 cavity.

For the simulations of the *apo* Octacid4 in water at 298 K, we found that the Rg of the cavity remained at ~5.4 Å and periodically spiked to >5.7 Å (Fig. 5A and Fig. S1), while the cavity Rgs of the open and closed conformations of Octacid4 in vacuo were 5.86 Å and 5.37 Å, respectively (Table 1). The average and standard error of the cavity Rg for the 20 simulations at 298 K were 5.44 Å and 0.02 Å, respectively (Table 1). For the simulations at 340 K, the Rg also remained at ~5.4 Å and spiked to >5.7 Å, but the frequency of the spikes was much higher at 340 K than that at 298 K (Fig. 5B and Fig. S1), and the average and standard error of the Rg at 340 K were 5.45 Å and 0.02 Å, respectively (Table 1).

**Fig. 5.**
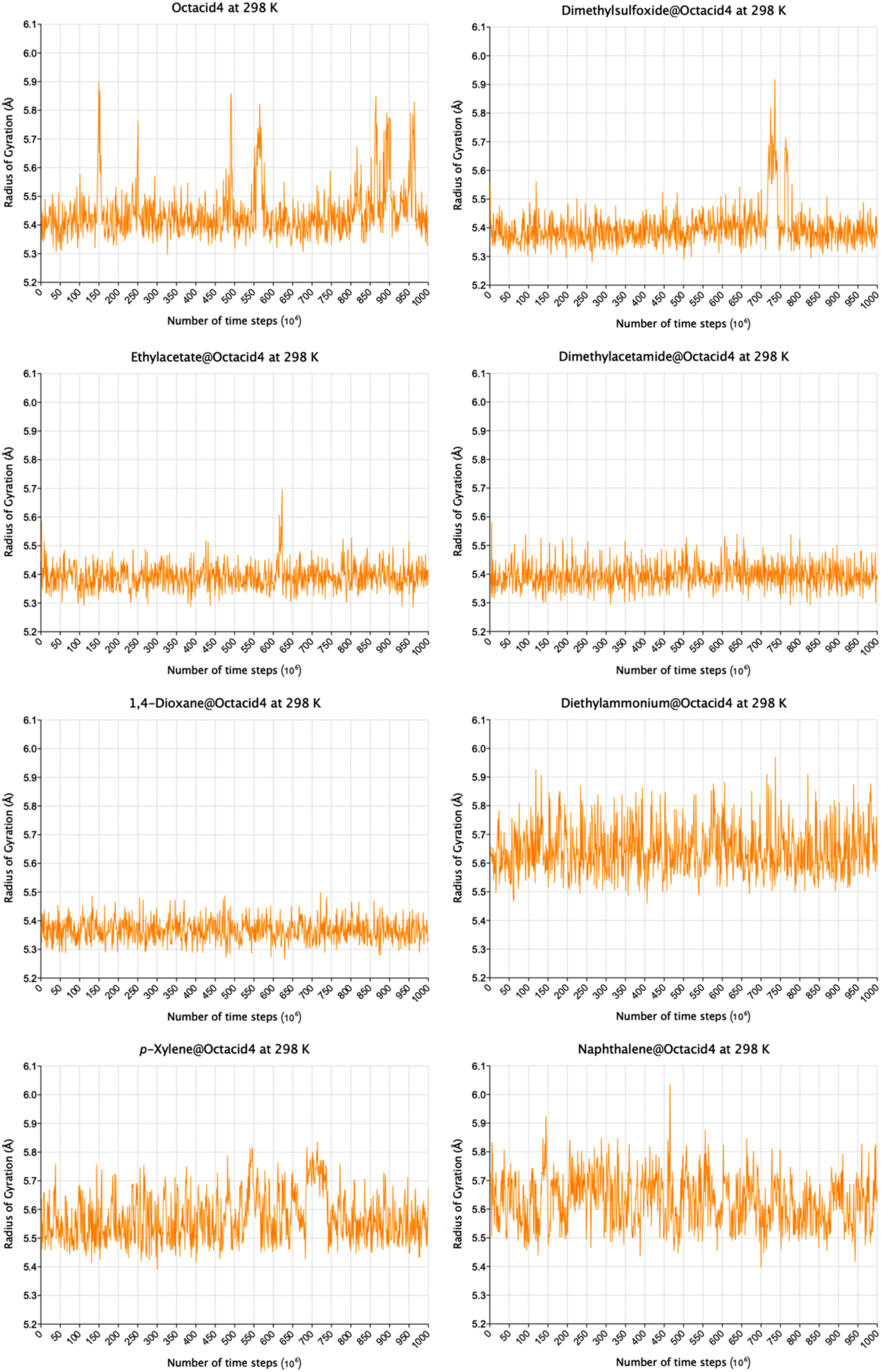
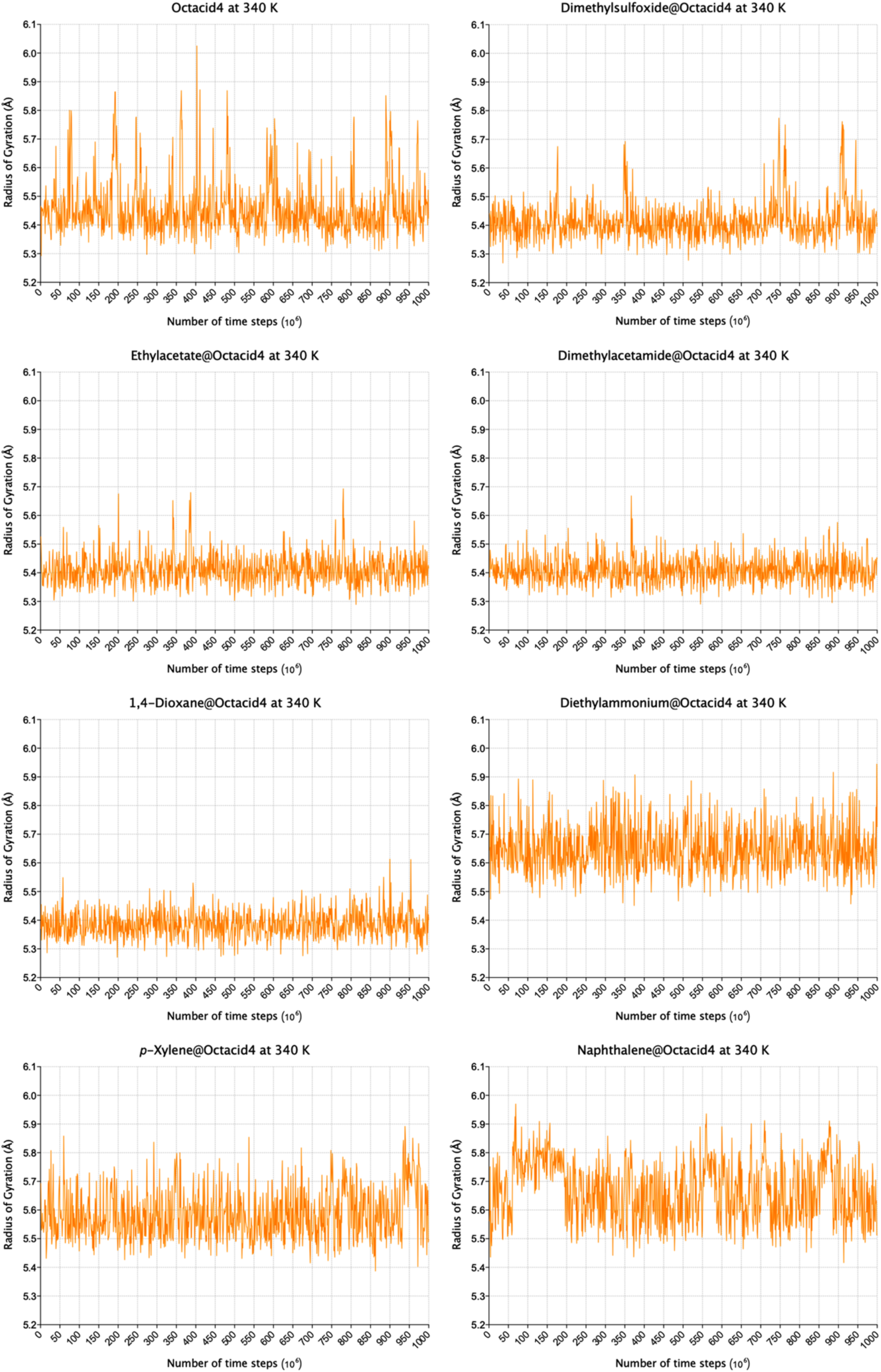
Time series of radius of gyration of the Octacid4 cavity for the first of 20 distinct and independent MD simulations. **(A)** Simulations at 298 K. **(B)** Simulations at 340 K.

These time series of Rg and their average values indicate that (1) *apo* Octacid4 in water adopted two clusters of conformations, one similar to the closed conformation of *apo* Octacid4 in vacuo with a small cavity and the other similar to the open conformation of *apo* Octacid4 in vacuo with a large cavity, (2) the large-cavity conformations were much less populated in water than the small-cavity conformations, (3) the population of the large-cavity conformations in water increased as temperature increased, and (4) *apo* Octacid4 in water underwent cavity expansion and contraction at both temperatures.

For the simulations of the DMSO, EtOAc, and DMA hemicarceplexes, we found a cavity Rg of ~5.4 Å with a few or no spikes of >5.7 Å at 298 K and 340 K (Fig. 5 and Fig. S1); the respective average and standard error of the cavity Rg of each set of the 20 MD simulations at 298 K were 5.41 Å and 0.02 Å for DMSO, 5.40 Å and 0.01 Å for EtOAc, and 5.40 Å and 0.01 Å for DMA (Table 1). The corresponding values at 340 K were 5.41 Å and 0.02 Å for DMSO, 5.41 Å and 0.01 Å for EtOAc, and 5.41 Å and 0.01 Å for DMA (Table 1). For the simulations of the 1,4-dioxane hemicarceplex, we found a cavity Rg of ~5.35 Å with no spikes at either temperature (Fig. 5 and Fig. S1); the respective average and standard error of the cavity Rg for the simulations at 298 K were 5.37 Å and 0.01 Å, and the corresponding values at 340 K were 5.38 Å and 0.01 Å (Table 1). These time series of Rg and their average values indicate that (1) the hemicarceplexes involving relatively compact guests in water all adopted one cluster of conformations that is similar to the closed conformation of *apo* Octacid4 in vacuo except for DMSO•Octacid4 (which adopted two clusters conformations that resemble the open and closed conformations of *apo* Octacid4 in vacuo, but the population of the open conformation of DMSO•Octacid4 was much lower than that of *apo* Octacid4 in water), and (2) these hemicarceplexes mostly kept their cavities contracted with all equatorial portals closed.

For the simulations of the DEA, *p*-xylene, and naphthalene hemicarceplexes, we found a cavity Rg of ~5.6 Å with a few or no spikes of >5.7 Å at 298 K and 340 K (Fig. 5 and Fig. S1); the respective average and standard error of the cavity Rg of the simulations at 298 K were 5.64 Å and 0.02 Å for DEA, 5.56 Å and 0.02 Å for *p*-xylene, and 5.65 Å and 0.02 Å for naphthalene. The corresponding values at 340 K were 5.65 Å and 0.02 Å for DEA, 5.59 Å and 0.02 Å for *p*-xylene, and 5.66 Å and 0.02 Å for naphthalene (Table 1). These time series of Rg and their average values indicate that (1) the hemicarceplexes involving relatively bulky guests in water all adopted one cluster of conformations with their cavities larger than that of the closed conformation of *apo* Octacid4 in vacuo and smaller than that of the open conformation of *apo* Octacid4 in vacuo, and (2) these hemicarceplexes mostly kept their cavities contracted with all equatorial portals closed.

### Characterization of guest motion in Octacid4

We also performed conformational cluster analyses on the simulations at 340 K for *apo* Octacid4 and the seven hemicarceplexes. All hemicarceplex simulations were performed using an initial conformation in which the host adopted the open conformation of *apo* Octacid4 in vacuo (Fig. 2A) and the guest was manually docked into the host cavity with its maximal dimension perpendicular to the axial axis (*viz*., the axis passing the two axial portals). Interestingly, as detailed below, in the aqueous MD simulations, all seven guests adopted a new orientation with their maximal dimensions parallel to the axial axis (Fig. 6). Similar results were obtained (Fig. S2) when the analyses were performed using the simulations at 298 K.

**Fig. 6.**
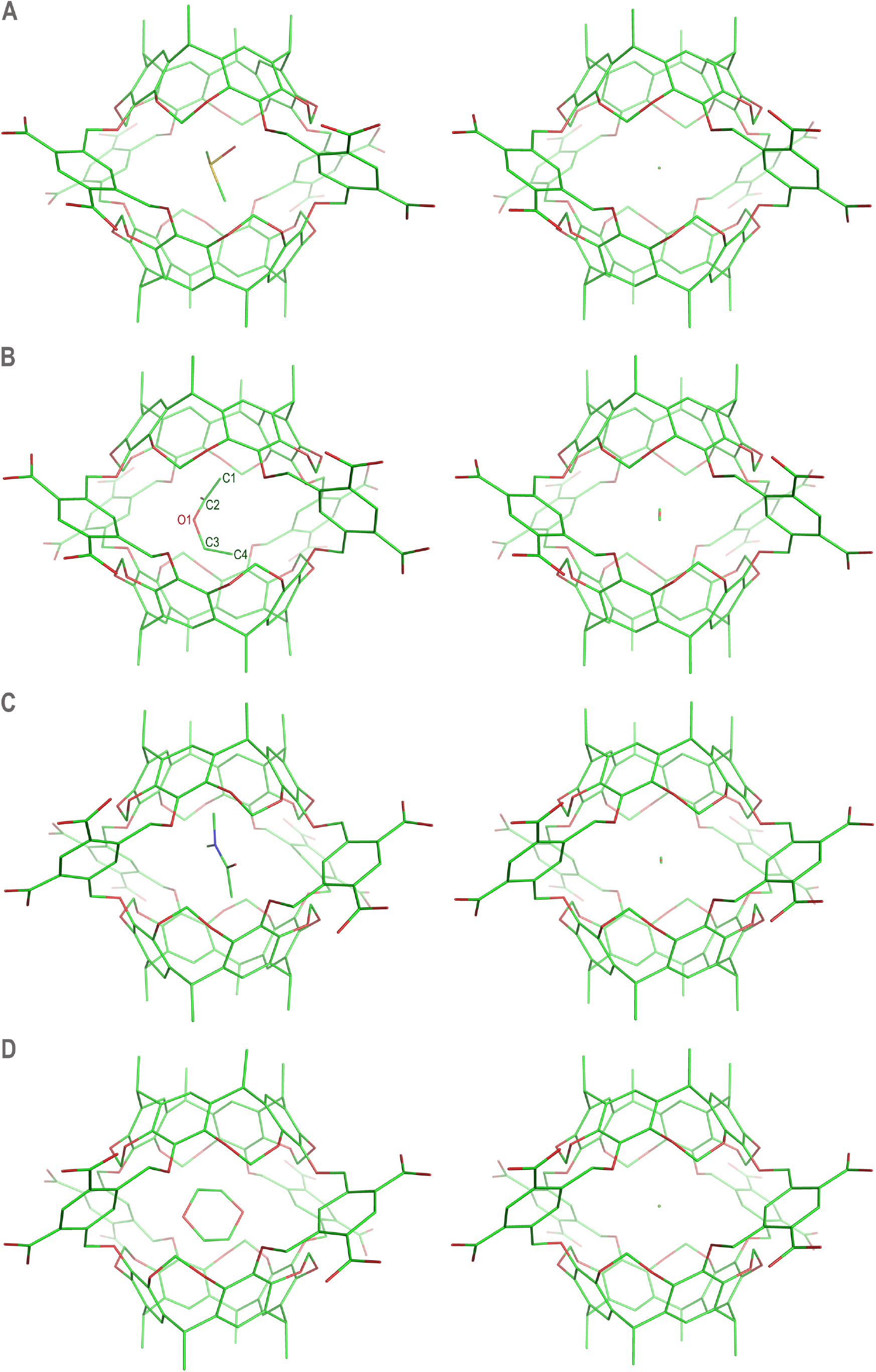
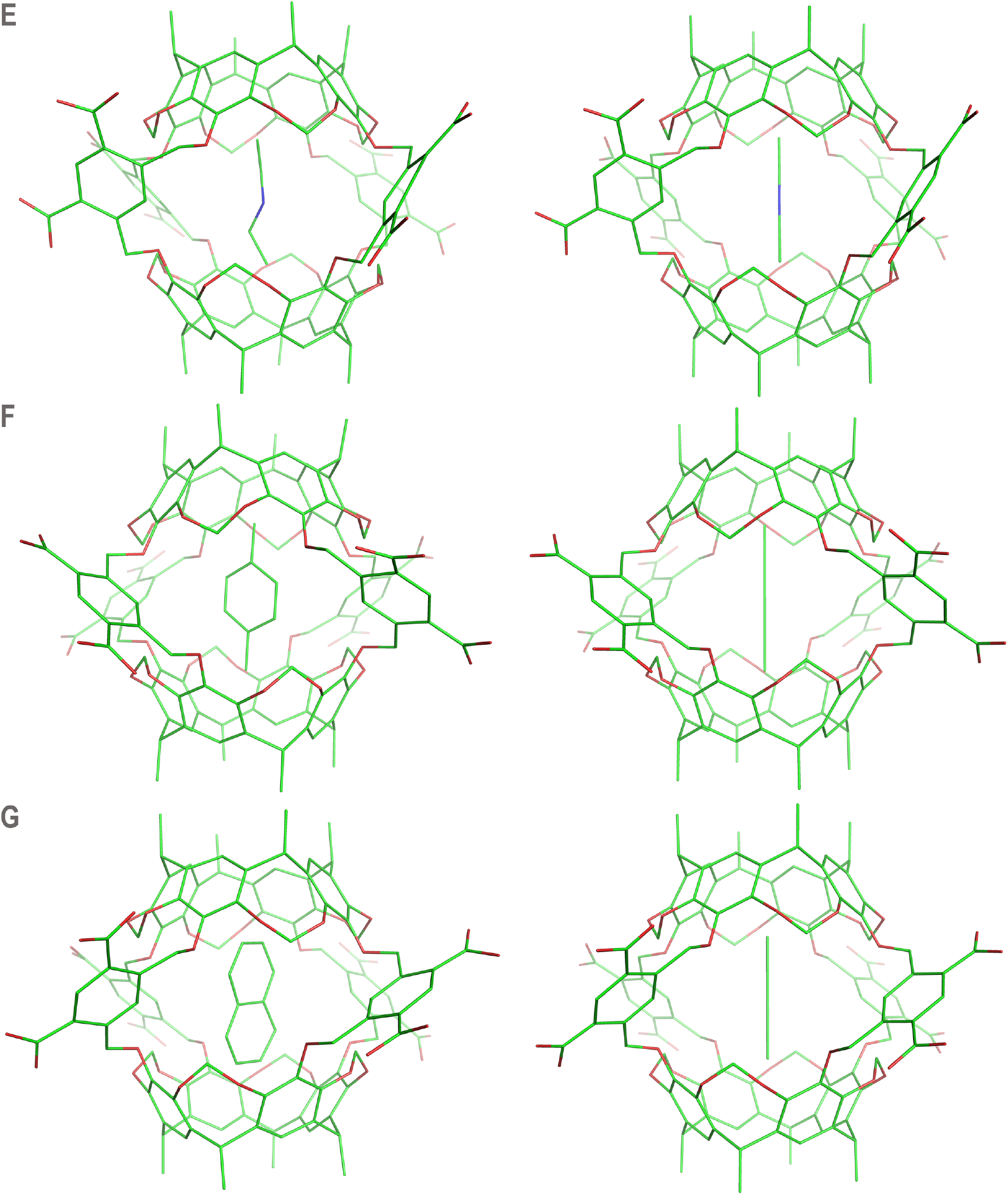
The most populated conformations of seven Octacid4 hemicarceplexes in the aqueous MD simulations at 340 K. The representative and average conformations in the largest conformation cluster of 20 MD simulations for each hemicarceplex are shown in the left and right panels, respectively. The sulfur, oxygen, nitrogen, and carbon atoms are in yellow, red, blue, and green, respectively. Counter ions and water molecules of aqueous Octacid4 are not displayed for clarity. **(A)** DMSO•Octacid4. **(B)** EtOAc•Octacid4. **(C)** DMA•Octacid4. **(D)**1,4-Dioxane•Octacid4. **(E)** DEA•Octacid4. **(F)***p*-Xylene•Octacid4. **(G)** Naphthalene•Octacid4.

For *apo* Octacid4 in water, the most populated conformation (occurrence: 97.2%) in the MD simulations was nearly identical to the closed conformation in vacuo (carbon root mean square deviation: 0.34 Å; Figs. 2A and 2C), whereas the second most populated conformation (occurrence: 1.9%) was somewhat different from the open conformation in vacuo (carbon root mean square deviation: 1.49 Å) in that the solution conformation had one equatorial portal open and the opposite portal closed (Figs. 2B and 2D).

For the DMSO hemicarceplex (Fig. 6A), the most populated conformation had the guest oxygen atom pointing to the equatorial portals and the two guest methyl groups pointing to the axial portals; in the average conformation of the largest conformation cluster, the guest was shrunk to a ball with the oxygen atom on one side and the two overlapping carbon atoms on the other, indicating that the guest had rotated around multiple axes.

For the EtOAc hemicarceplex (Fig. 6B), the most populated guest conformation (population: 94.7%) was not fully extended (with C1-C2-O1-C3 and C2-O1-C3-C4 being –114° and 81°, respectively), and the population of the fully extended guest conformation with corresponding torsions of –180° and 180° was 0.1%; the most populated guest conformation had its two oxygen atoms pointing to the equatorial portals and its two methyl groups pointing to the axial portals; in the average conformation of the largest conformation cluster, the guest was shrunk to a short rod with the oxygen atoms in the middle and the two carbon atoms on both ends of the rod, indicating that the guest had rotated frequently around the axial axis and less frequently around the equatorial axis (the axis passing the two opposing equatorial portals).

For the DMA hemicarceplex (Fig. 6C), the most populated conformation had the guest oxygen atom pointing to the equatorial portals and the two trans methyl groups of the guest pointing to the axial portals; in the average conformation of the largest conformation cluster, the guest was shrunk to a short rod with (1) the oxygen atom on one side of the middle region, (2) the carbon atom trans to the oxygen atom on the other side of the middle region, and (3) the two trans carbon atoms on both ends of the rod, indicating that the guest had rotated frequently around the axial axis and less frequently around the equatorial axis.

The 1,4-dioxane hemicarceplex (Fig. 6D) had one conformation cluster. The most populated guest conformation adopted the energetically stable chair conformation with the two guest oxygen atoms pointing to the equatorial portals and the two methylene groups on one side of the guest pointing to an axial portal while those on the other side pointing to the opposite axial portal. In the average conformation of the cluster, the guest was shrunk to a ball with two oxygen atoms on opposite sides, indicating that the guest had rotated around multiple axes.

The DEA, *p*-xylene, and naphthalene hemicarceplexes (Figs. 6E–6G) had one conformation cluster each. The conformations of these clusters had (1) the DEA nitrogen atom pointing to the equatorial portals and the two DEA methyl groups pointing to the axial portals, (2) the two *p*-xylene methyl groups pointing to the axial portals, and (3) the naphthalene β-carbon atoms pointing to the axial portals. In the average conformation of each cluster, the guest was shrunk to a long rod whose length was the same as that of the guest, indicating that these guests all had rotated exclusively around the axial axis.

## Discussion

### Support for the modular approach

Our characterization studies reveal that (1) *apo* Octacid4 adopted two clusters of conformations in aqueous MD simulations at 298 K, (2) one cluster had the conformations (population: 97.2%) nearly identical to the one in vacuo with all equatorial portals closed, and (3) the other had the conformations (population: 1.9%) somewhat different from the open conformation of *apo* Octacid4 in vacuo in that the solution conformation has one equatorial portal open and the opposite portal closed. The population ratio (97.2/1.9) of the two clusters according to the conformational cluster analysis is consistent with the frequency for the conformational exchanges depicted in Fig. 5, which was derived from the Rg-based analysis. In addition, all solution conformations derived from the MD simulations have one resorcinarene fragment rotated by ~22° with respect to the other, which is consistent with reported crystal structures of different hemicarceplexes (involving the same or different guests) all of which have one resorcinarene fragment rotated by 13–21° with respect to the other^1,4,19,20^. Moreover, the less populated aqueous Octacid4 conformations with one equatorial portal open and the opposite portal closed are also consistent with the reported V-shaped hemicarcerand conformation proposed in the sliding-door mechanism for hemicarceplex formation^7,21^. These internal and external consistencies indicate the plausibility of the aqueous conformations of Octacid4 obtained a priori from our modular-approach– based MD simulations and lend credence to our modular approach to hemicarcerand simulations for probing solution conformations relevant to the following mechanistic insights.

### How Octacid_4_ incarcerates its guests

Our characterization studies also show that *apo* Octacid4 in water mostly adopted a cluster of conformations with small cavities whose equatorial portals are all closed and that, even at 298 K, this host also periodically and transiently adopted another cluster of conformations that have large cavities with at least one equatorial portal open. These periodic and transient conformations with an open equatorial portal explain how Octacid4 reportedly encapsulated guests in only a few minutes at 298 K in the NMR experiments^3^, but do not explain how Octacid4 reportedly kept its guests encapsulated at the same temperature^3^. Perhaps the guests were incarcerated by strong intrinsic binding (*viz*., the complexation governed by strong nonbonded intermolecular interactions). However, the involvement of strong intrinsic binding is debatable given the weak interaction of the cavity with DMSO or 1,4-dioxane as indicated by their free spins as described above (Table 1 and Figs. 6A and 6D). An alternative explanation is therefore needed.

Unexpectedly, our characterization studies show that upon complexation, Octacid4 adopted only one cluster of conformations. These conformations have, depending on the size of the guest, a contracted or expanded cavity relative to that of *apo* Octacid4 in water. The conformations with contracted cavities have the equatorial portals all closed because the guests are relatively compact and subsequently the weak nonbonded intermolecular interactions of the host cavity with a small guest shift the host conformations with an open equatorial portal to the conformations with all equatorial portals closed. The conformations with slightly expanded cavities also have the equatorial portals all closed because the guests are relatively bulky and these guests jam the cavity and consequently prevent the host from adopting the V-shaped sliding-door conformations shown in Figs. 2D and 2E to open any of the equatorial portals. Therefore, regardless of the size of the guest, the guest-bound Octacid4 in water adopts one cluster of conformations with closed equatorial portals. These conformations satisfactorily explain how Octacid4 can incarcerate a range of guests at 298 K, which is the temperature at which the guests enter the cavity.

### Implication

Collectively, our studies suggest that Octacid4 can open one of its equatorial portals without elevating temperature before complexation and close all of its equatorial portals without lowering temperature after complexation. These interesting and unexpected capabilities enable the unique constrictive binding of Octacid4 with a range of small-molecule guests and suggest further that the guest-induced host conformational change that impedes decomplexation is a previously unrecognized conformational characteristic that is conducive to strong molecular complexation. This characteristic could broaden the theoretical dissection of the experimentally observed complexation affinity and the design of new complex systems for materials technology, data storage and processing, catalysis, drug design and delivery, and medicine by accounting for not only intrinsic binding (which is limited because, confined by the cost of chemical synthesis, only a small number of functional groups can be introduced to improve nonbonded intermolecular interactions) but also constrictive binding (which requires one or a few functional groups to trigger the formation of a host conformation that hinders decomplexation as demonstrated by the Octacid4 hemicarceplexes).

## Methods

### Molecular dynamics simulations

Hemicarcerand Octacid4 or its hemicarceplex neutralized with sodium ions was solvated with the TIP3P water^13^ (“solvatebox molecule TIP3BOX 8.2”) using tLEaP of the AmberTools 16 package (University of California, San Francisco) and then energy-minimized for 100 cycles of steepest-descent minimization followed by 900 cycles of conjugate-gradient minimization to remove close van der Waals contacts using SANDER of the AMBER 11 package (University of California, San Francisco), forcefield FF12MClm^12^, and a cutoff of 8.0 Å for nonbonded interactions. The resulting system was heated from 5 K to 298 or 340 K at a rate of 10 K/ps under constant temperature and constant volume, and then equilibrated for 10^6^ timesteps under constant temperature of 298 K or 340 K and constant pressure of 1 atm employing isotropic molecule-based scaling. Finally, a set of 20 distinct, independent, unrestricted, unbiased, isobaric–isothermal, and 316-ns MD simulations of the equilibrated system was performed for Octacid4 or its hemicarceplex using PMEMD of the AMBER 16 package (University of California, San Francisco), forcefield FF12MClm^12^, and a periodic boundary condition at 298 K or 340 K and 1 atm. The 20 unique seed numbers for initial velocities of the 20 simulations were taken from Ref. ^22^. All simulations used (*i*) a dielectric constant of 1.0, (*ii*) the Berendsen coupling algorithm^23^, (*iii*) the particle mesh Ewald method to calculate electrostatic interactions of two atoms at a separation of >8 Å^24^, (*iv*) ∆*t* = 1.00 fs of the standard-mass time^12,25^, (*v*) the SHAKE-bond-length constraint applied to all bonds involving hydrogen, (*vi*) a protocol to save the image closest to the middle of the “primary box” to the restart and trajectory files, (*vii*) a formatted restart file, (*viii*) the revised alkali ion parameters^26^, (*ix*) a cutoff of 8.0 Å for nonbonded interactions, (*x*) a uniform 10-fold reduction in the atomic masses of the entire simulation system (both solute and solvent)12,25, and (*xi*) default values of all other inputs of PMEMD. The *apo* Octacid4 conformation with all equatorial portals open (Fig. 2B) was used as the initial conformation for the *apo* Octacid4 MD simulations. This conformation was obtained from energy minimization for 200 cycles of steepest-descent minimization followed by 2800 cycles of conjugate-gradient minimization using SANDER, forcefield FF12MClm^12^, and a cutoff of 30.0 Å for nonbonded interactions. For each hemicarceplex, the guest was manually docked into the cavity of the *apo* Octacid4 conformation with all equatorial portals open (Fig. 2B) in such a way that the maximal dimension of the guest was perpendicular to the axial axis (*viz.*, the axis passing the two axial portals); the resulting hemicarceplex was then energy minimized using the same minimization protocol for obtaining the *apo* Octacid4 conformation; the energy-minimized hemicarceplex was finally used as the initial conformation for the hemicarceplex MD simulations.

The forcefield parameters for the Octacid4 building block and for all small-molecule guests except dimethyl sulfoxide were obtained from ab initio calculations of the molecules at the HF/6-31G*//HF/6-31G* level using (1) a published procedure for the charge derivation^16^ and (2) the arithmetic average from multiple conformations for the bond, angle, and torsion parameters. The forcefield parameters for dimethyl sulfoxide were obtained from Ref. ^27^. The forcefield parameters of FF12MClm are available in the Supporting Information of Ref. ^25^.

All simulations were performed using computers at the University of Minnesota Supercomputing Institute and the Mayo Clinic high performance computing facility at the University of Illinois Urbana-Champaign National Center for Supercomputing Applications.

### ab Initio calculations

All ab initio calculations were performed using Gaussian 98 (Revision A.7; Gaussian, Inc. Wallingford, CT), except those for the intact Octacid4 at the HF/6-31G*//HF/6-31G* level were done using Gaussian 16 (Revision C.01; Gaussian, Inc. Wallingford, CT).

### Conformational cluster analysis

The conformational cluster analyses were performed using CPPTRAJ of the AmberTools 16 package (University of California, San Francisco) with the average-linkage algorithm^28^, epsilon of 1.0 Å, and root mean square coordinate deviation on all carbon atoms of Octacid4 or its hemicarceplex. Centering the coordinates of the hemicarceplex and imaging the coordinates to the primary unit cell were performed prior to the cluster analyses.

### Root mean square deviation and radius of gyration

Carbon root mean square deviations were calculated manually using ProFit V2.6 (http://www.bioinf.org.uk/software/profit/). Radius of gyration was calculated using CPPTRAJ.

## Data availability

All data generated for this study are included in this Article and its Supplementary Information.

## Acknowledgements

This work was supported by the US Army Research Office (W911NF-16-1-0264) and the Mayo Foundation for Medical Education and Research. Responsibility for the information and views in this study lies entirely with the authors. The authors acknowledge the computing resources provided by the University of Minnesota Supercomputing Institute and the Mayo Clinic high performance computing facility at the University of Illinois Urbana-Champaign National Center for Supercomputing Applications.

## Author contributions

K.G.M. performed the literature search; prepared the z-matrices of the HC1 and HCD1 residues; performed energy minimization of HC1, HCD1, and Octacid4; analyzed the MD simulation result of *p*-xylene•Octacid4; contributed to revisions of the manuscript. Y.-P.P. conceived the modular method; designed the HC1 and HCD1 residues and all protocols for MD simulation and analysis; performed the remaining computational study; analyzed the data; wrote the manuscript.

## Supplementary information

**Fig. S1. Time series of radius of gyration of the Octacid4 cavity for all 20 distinct and independent MD simulations at 298 K or 340 K.**

**Fig. S2. The most populated conformations of seven Octacid4 hemicarceplexes in the aqueous MD simulations at 298 K.** The representative and average conformations in the largest conformation cluster of 20 MD simulations for each hemicarceplex are shown in the left and right panels, respectively. The sulfur, oxygen, nitrogen, and carbon atoms are in yellow, red, blue, and green, respectively. Counter ions and water molecules of aqueous Octacid4 are not displayed for clarity. (A) DMSO•Octacid4. **(B)** EtOAc•Octacid4. **(C)** DMA•Octacid4. **(D)**1,4-Dioxane•Octacid4. **(E)** DEA•Octacid4. **(F)***p*-Xylene•Octacid4. **(G)** Naphthalene•Octacid4.

