## Supplementary figures and images for "How a Hemicarcerand Incarcerates Guests at Room Temperature Decoded with Modular Simulations"

### Supplemental Fig. S1

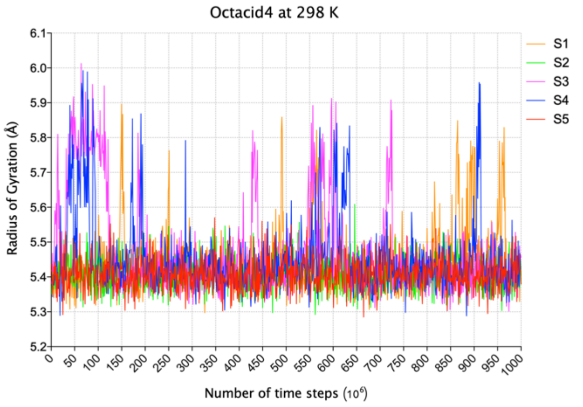

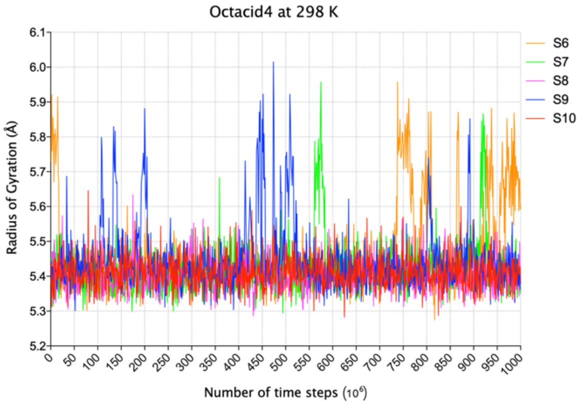

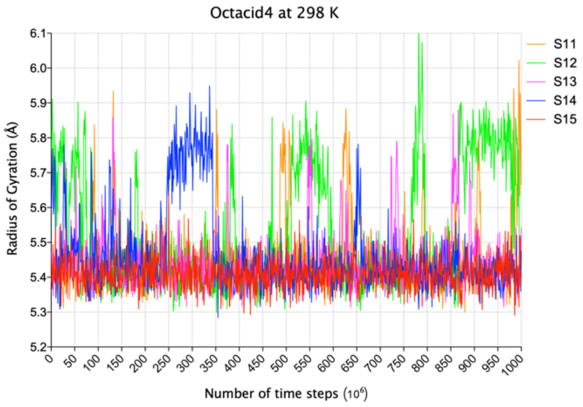

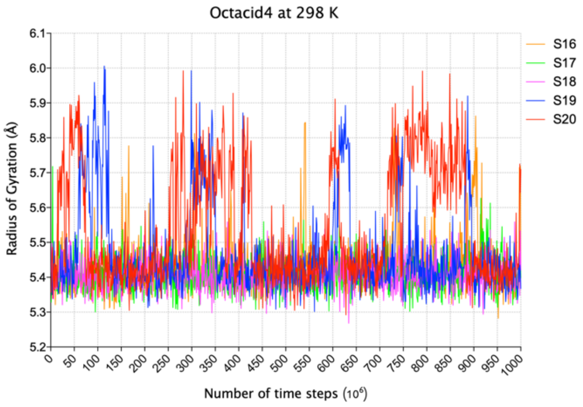

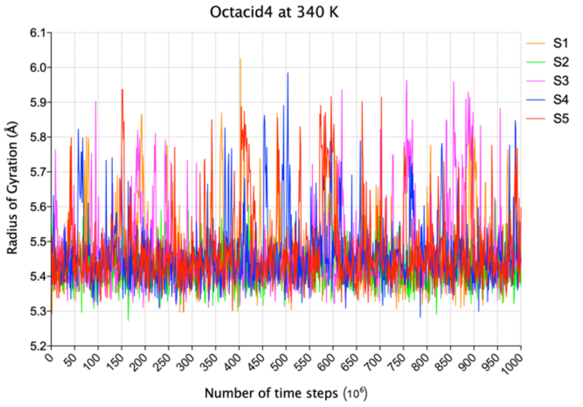

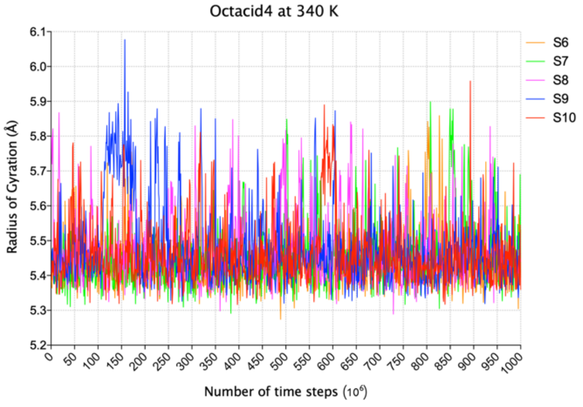

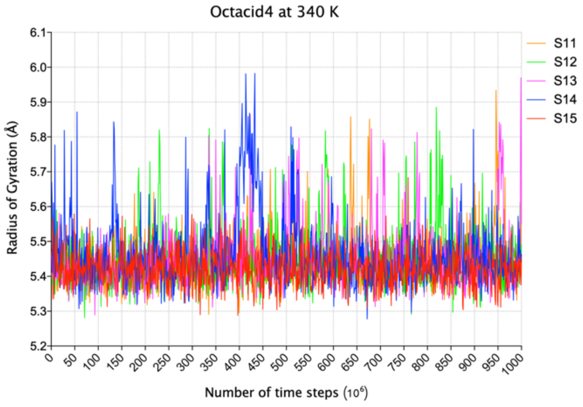

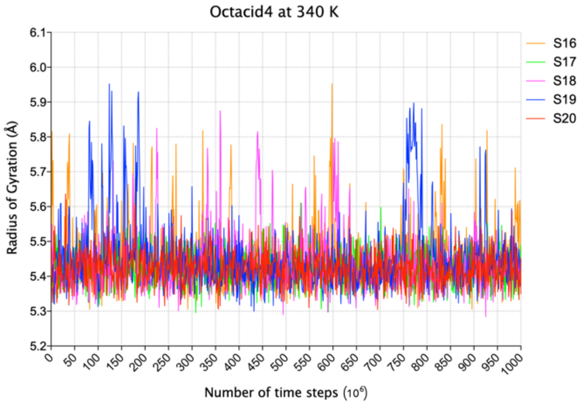

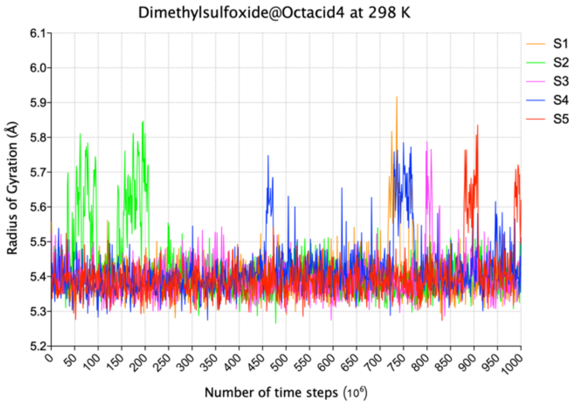

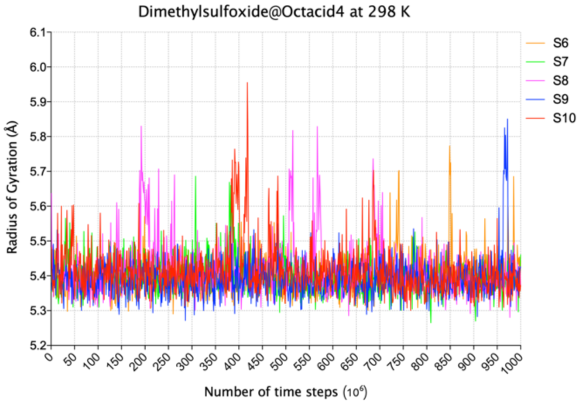

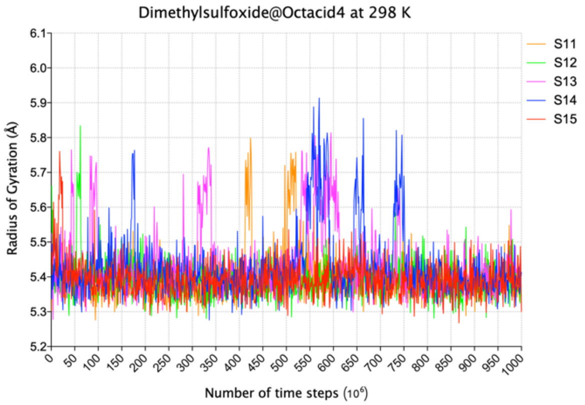

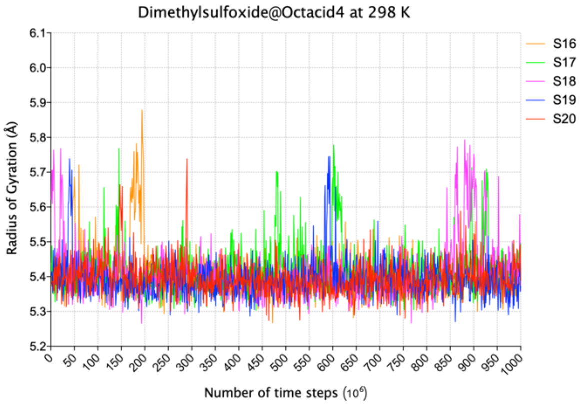

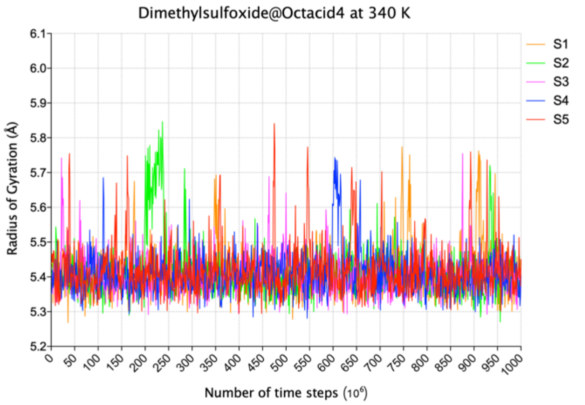

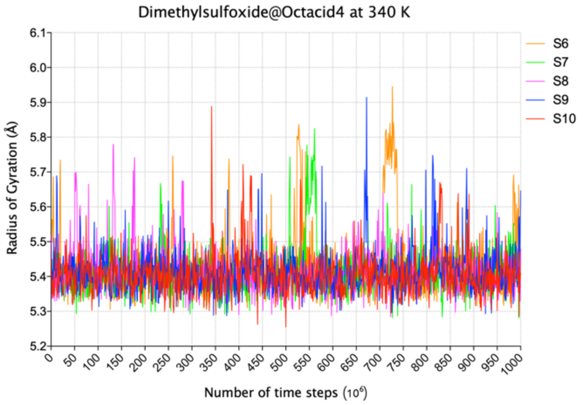

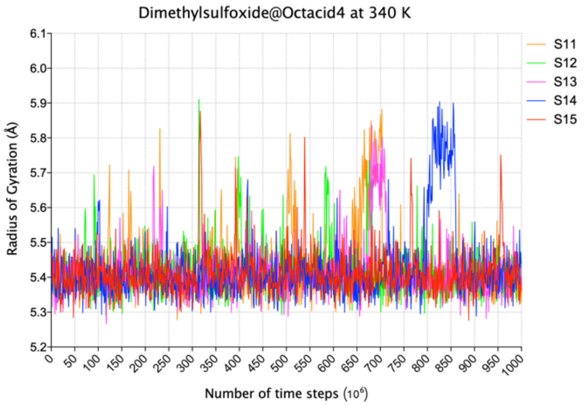

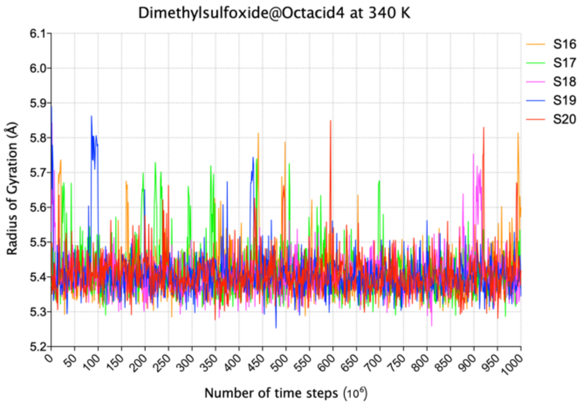

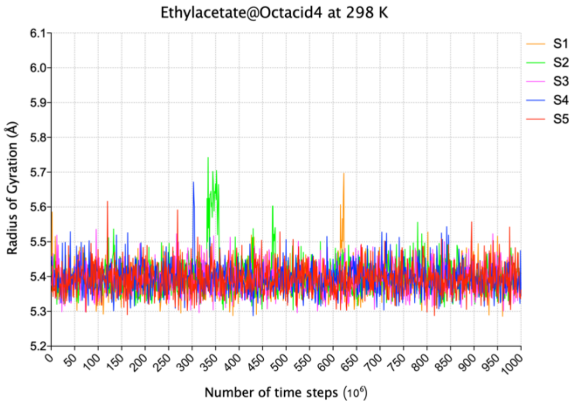

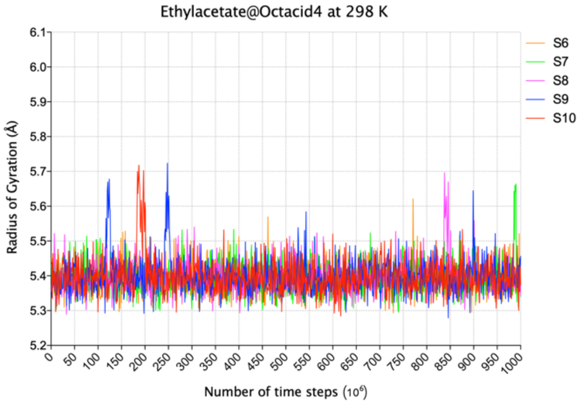

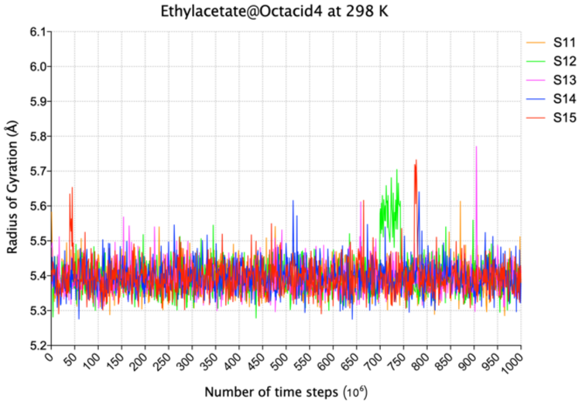

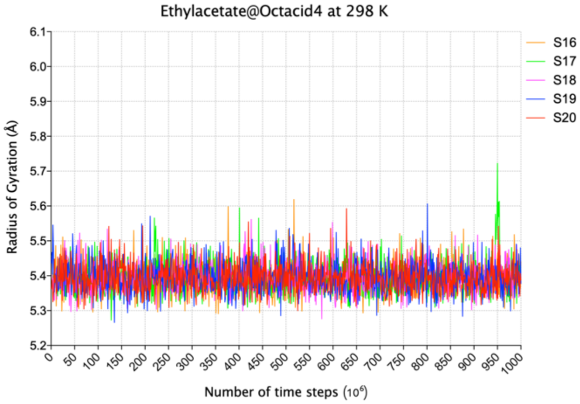

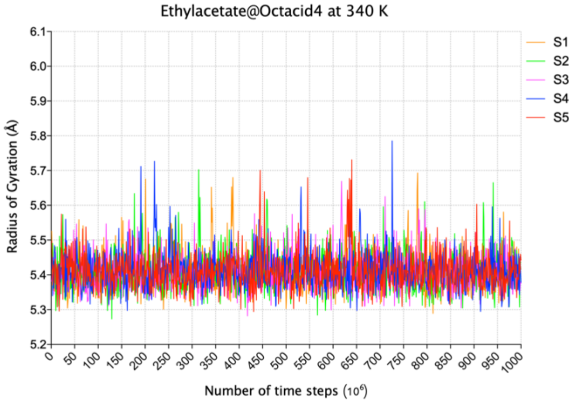

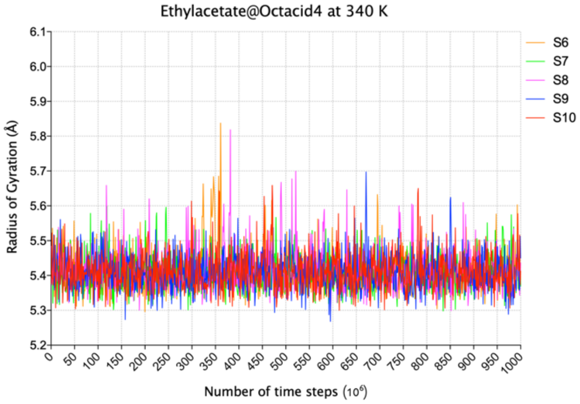

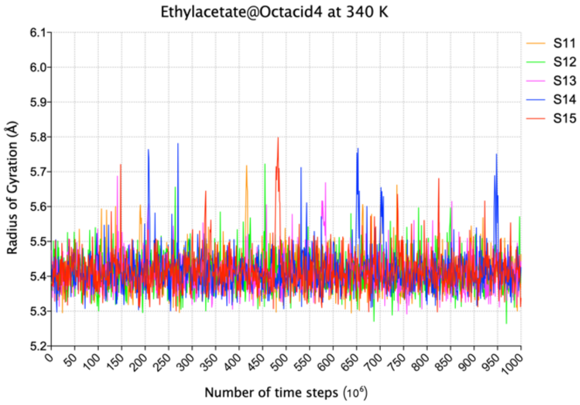

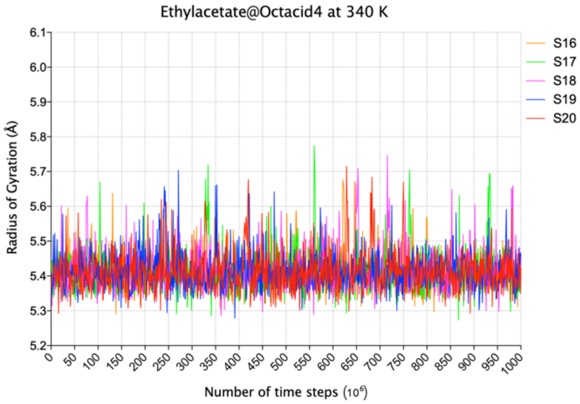

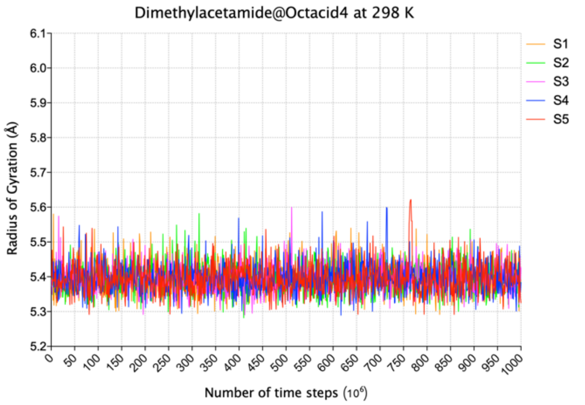

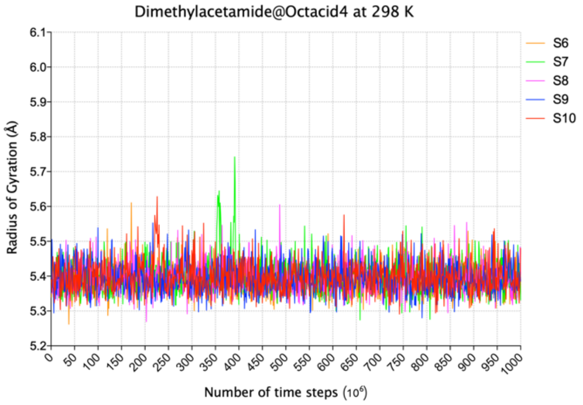

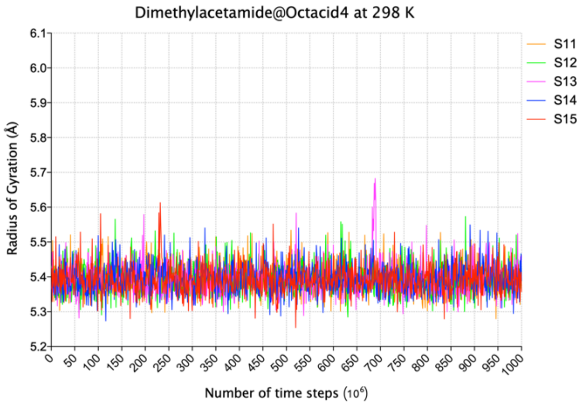

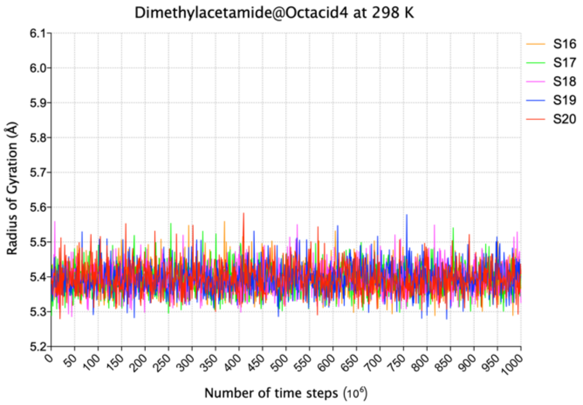

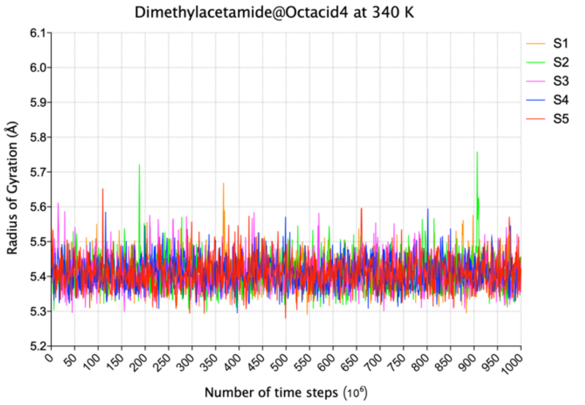

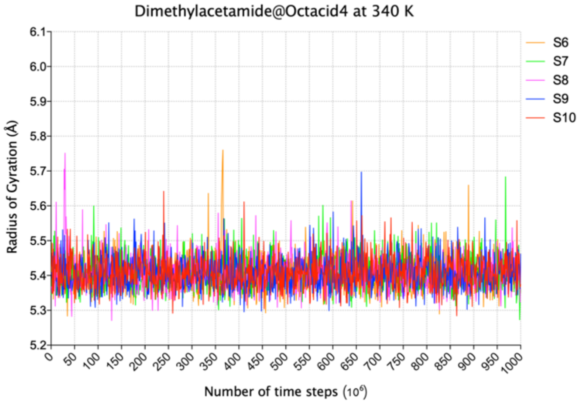

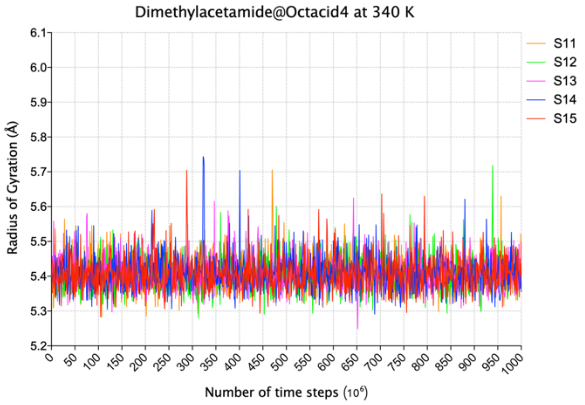

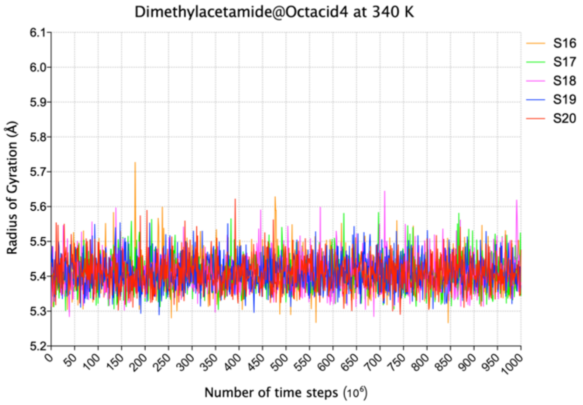

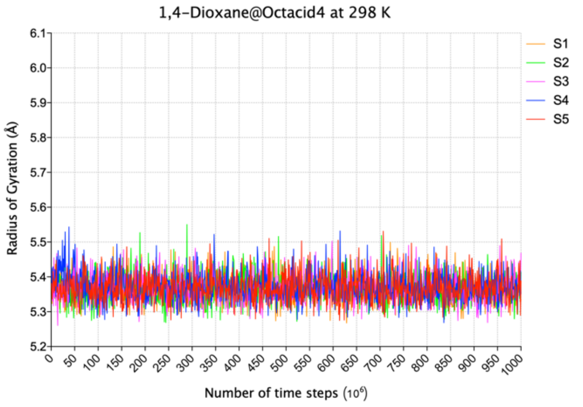

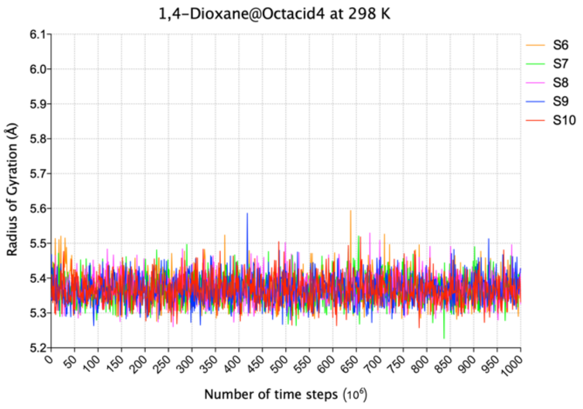

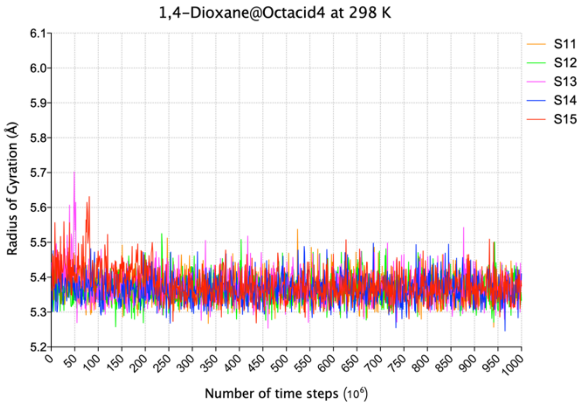

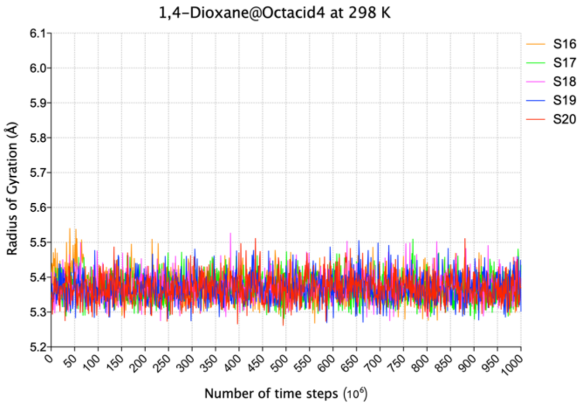

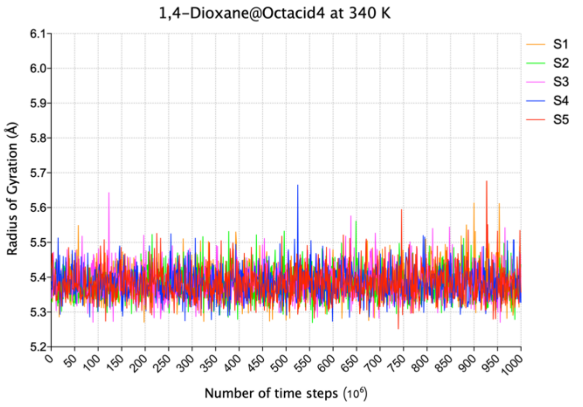

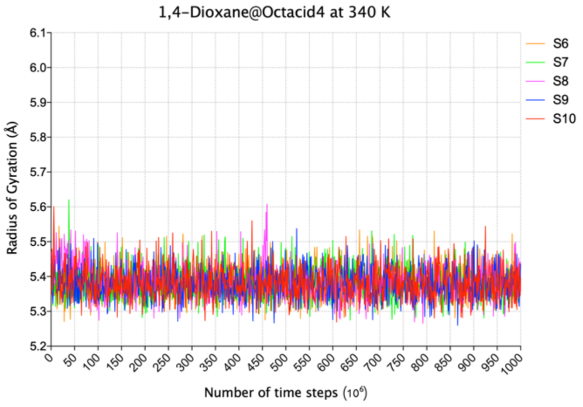

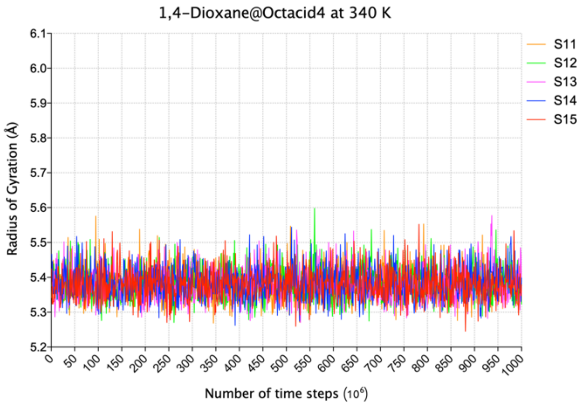

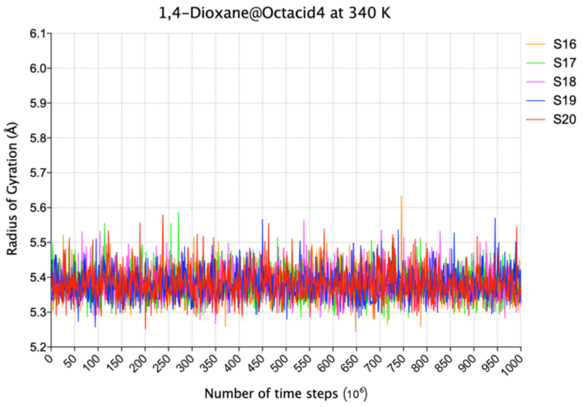

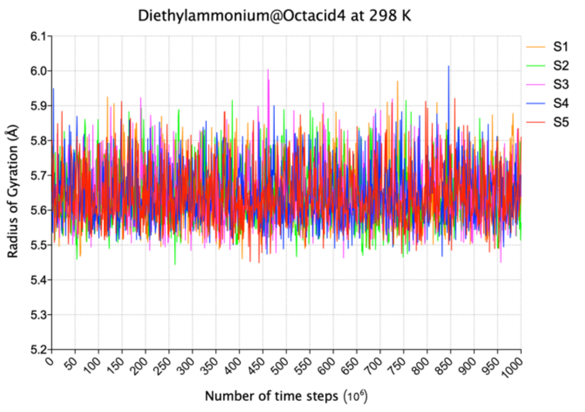

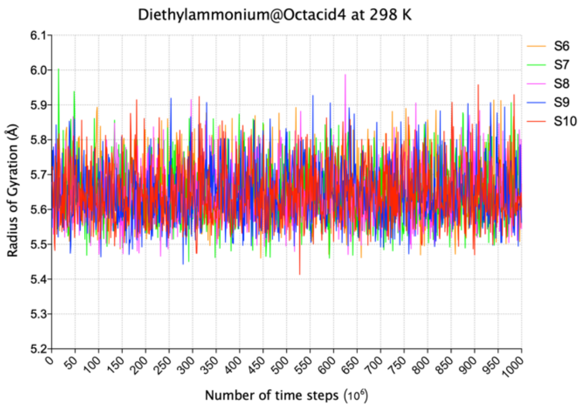

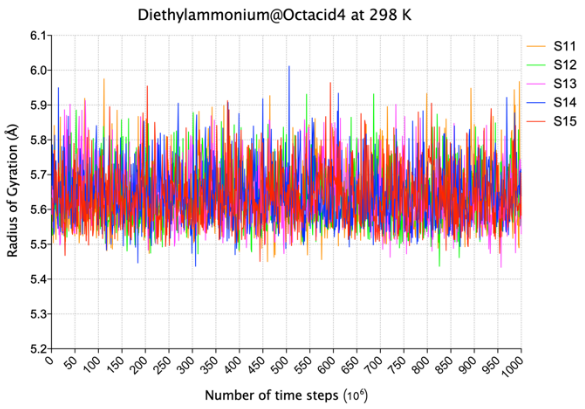

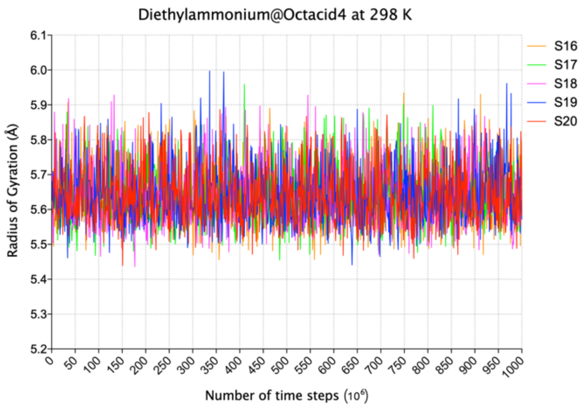

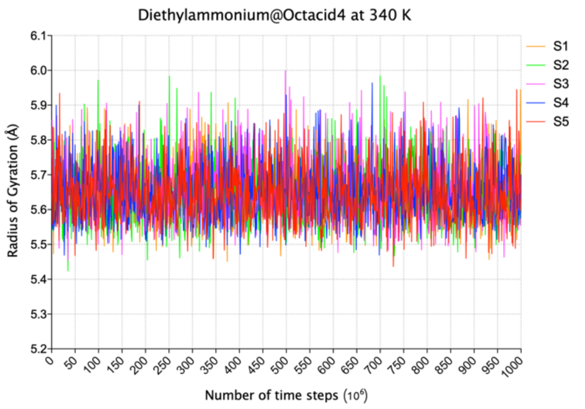

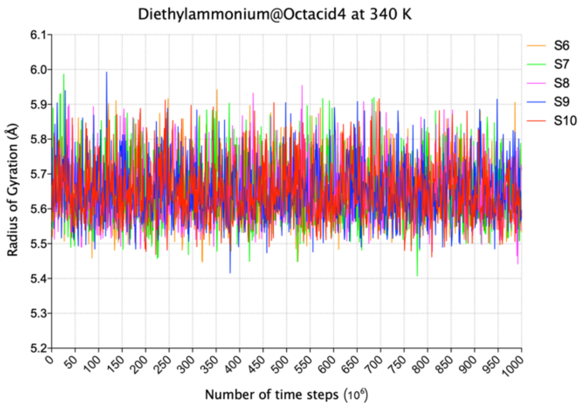

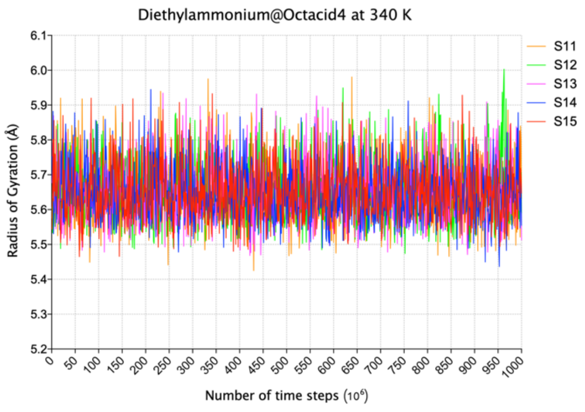

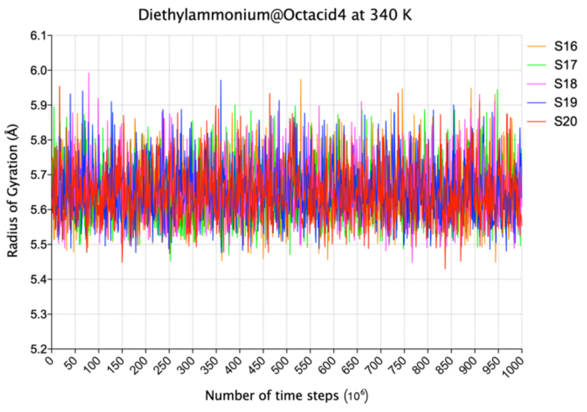

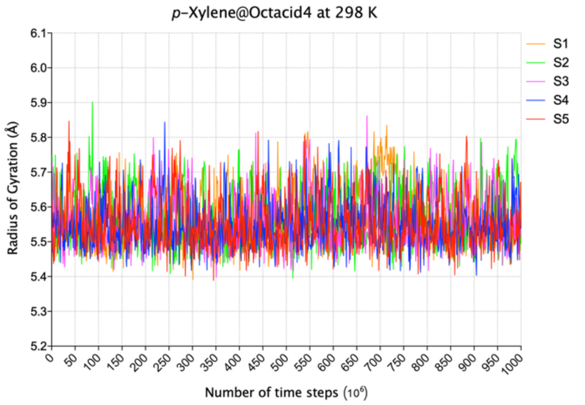

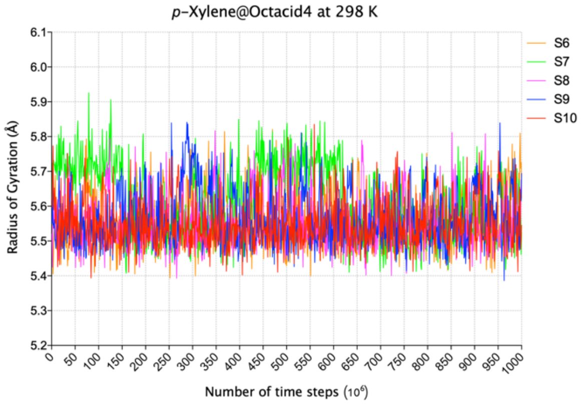

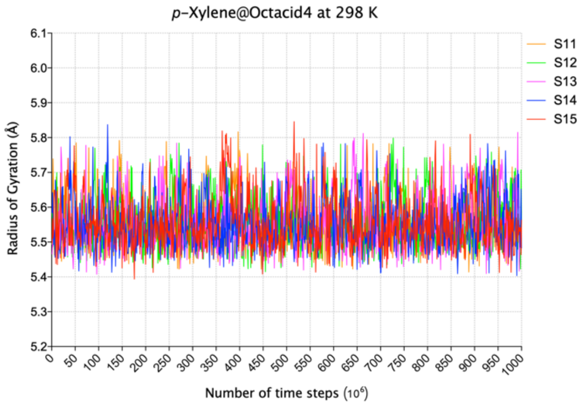

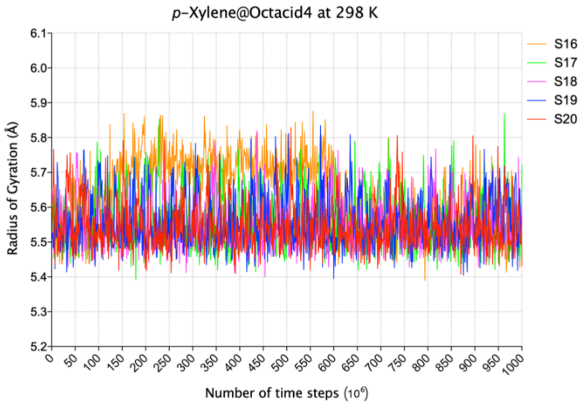

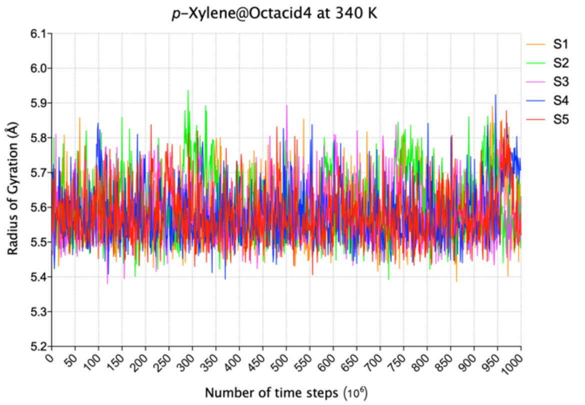

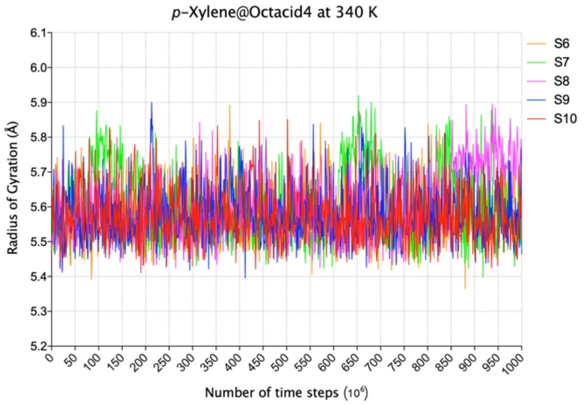

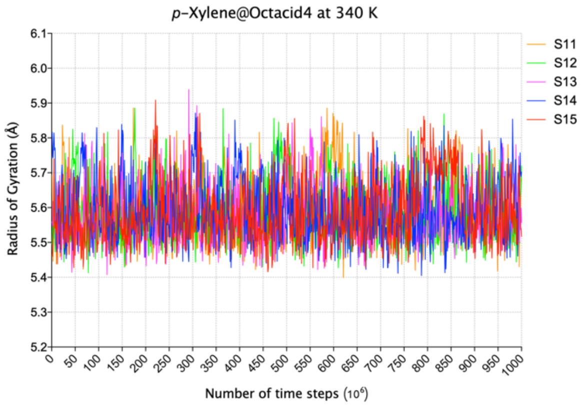

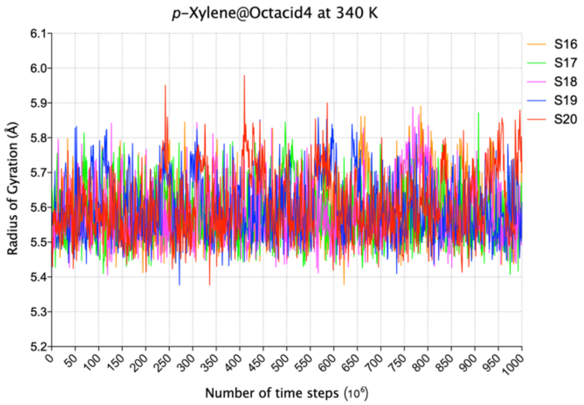

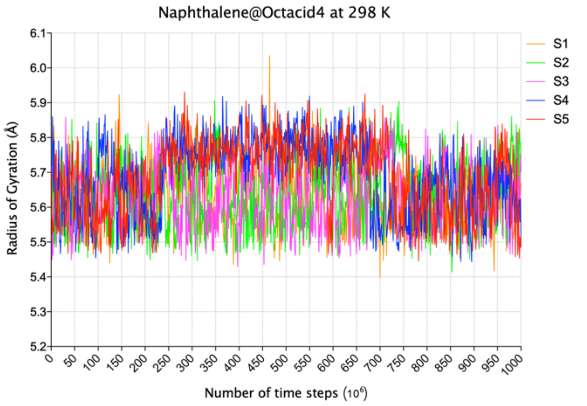

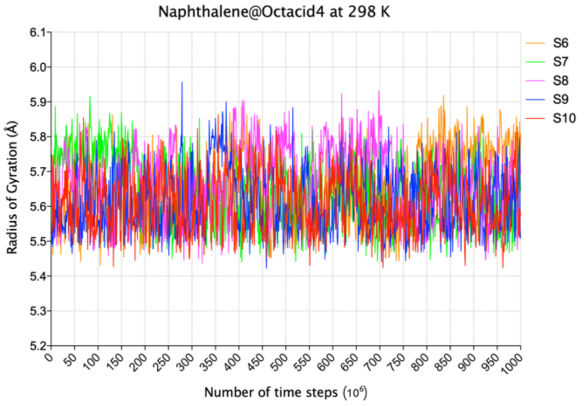

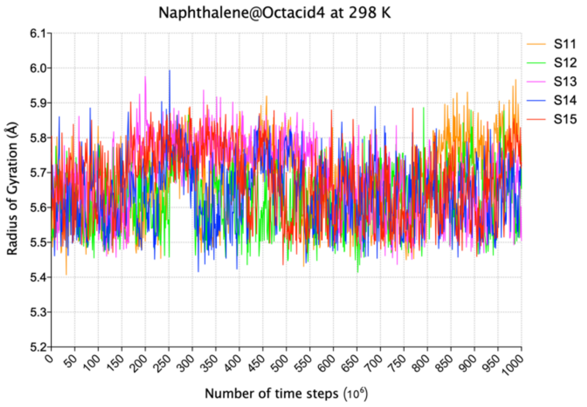

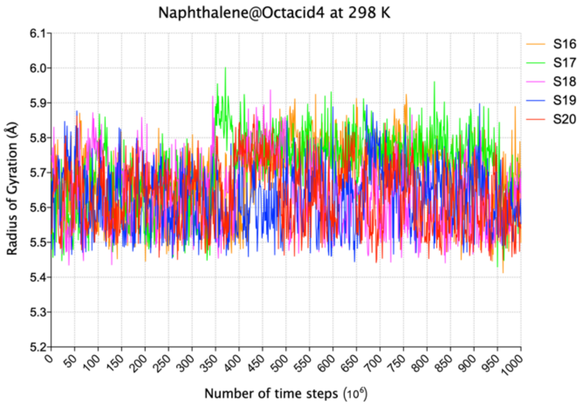

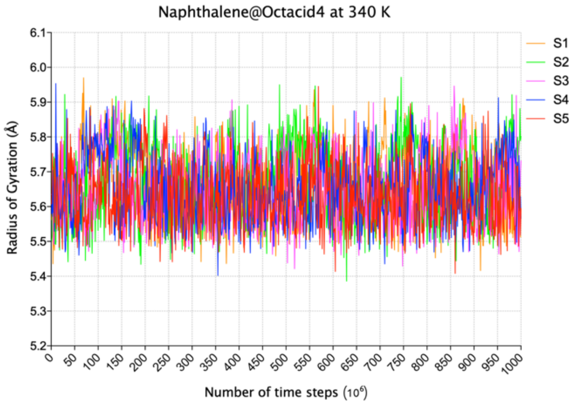

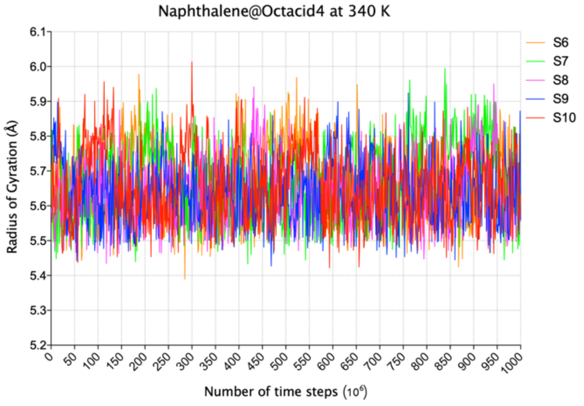

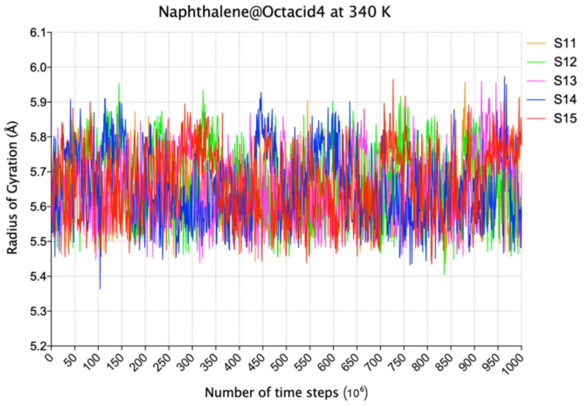

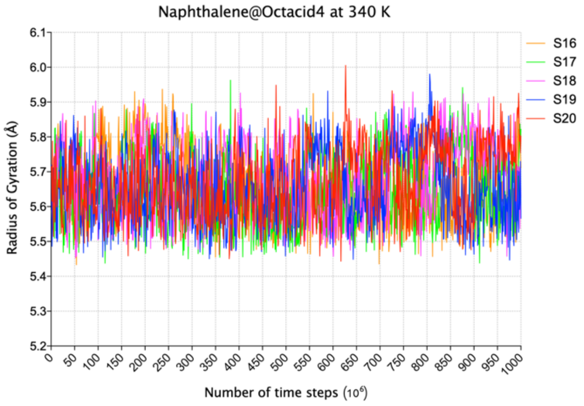

### Supplemental Fig. S2

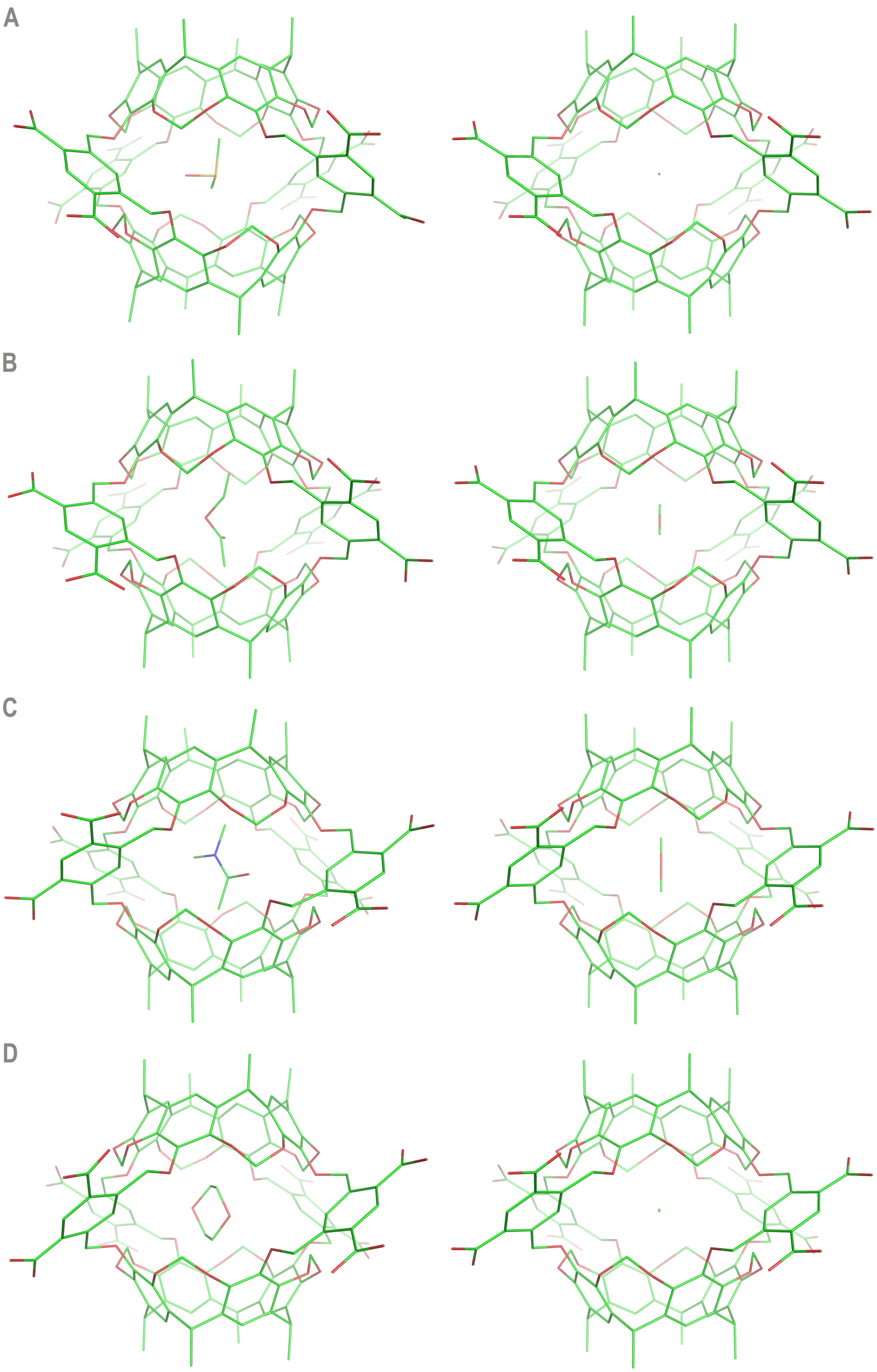

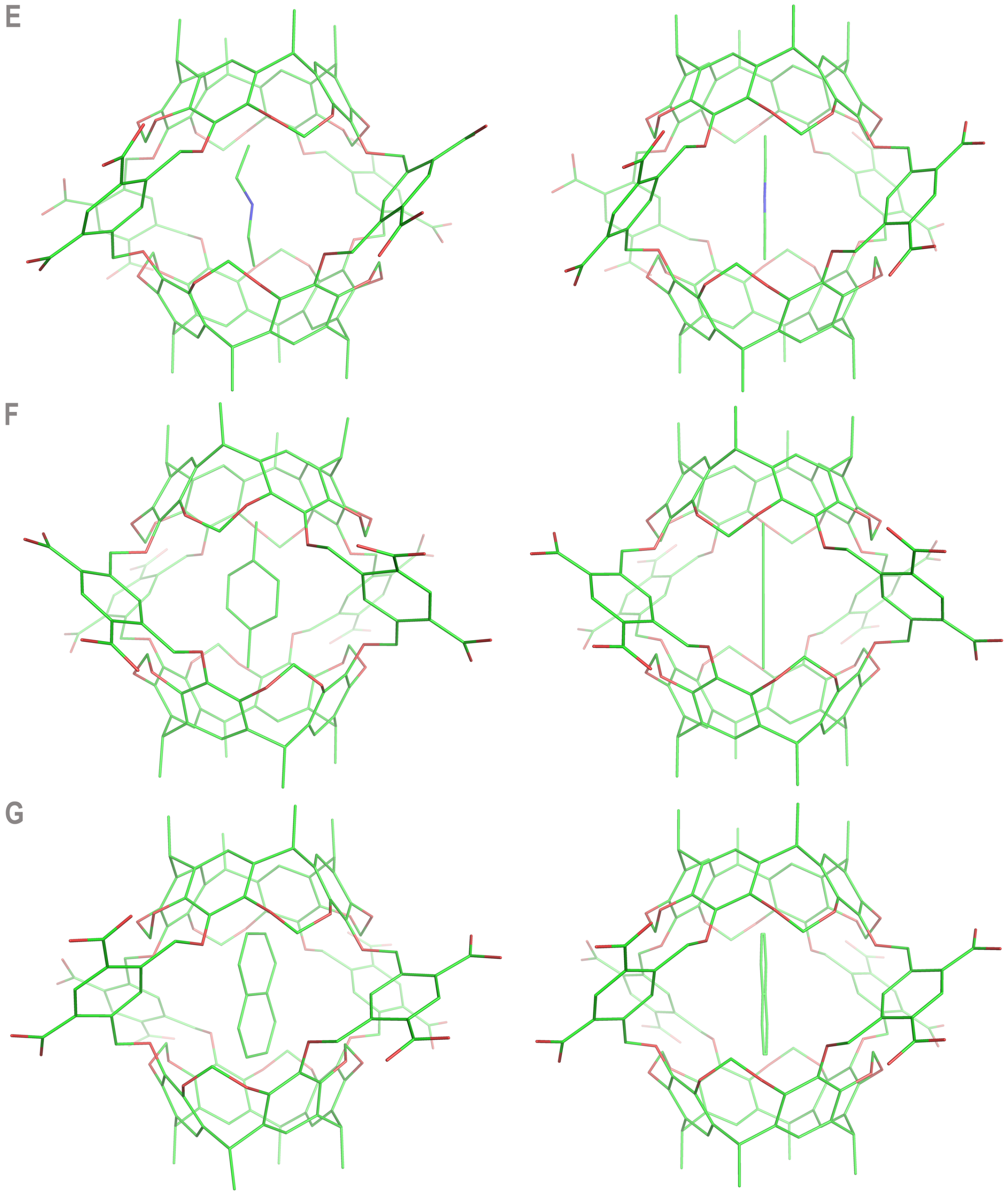
